# Phylogenomics of the North American Desert Radiation *Linanthus* (Polemoniaceae) Reveals Mixed Trait Lability and No Single Geographic Mode of Speciation

**DOI:** 10.1101/2024.06.13.598867

**Authors:** Ioana G. Anghel, Lydia L. Smith, Isaac H. Lichter-Marck, Felipe Zapata

**Affiliations:** Department of Ecology and Evolutionary Biology, University of California, Los Angeles, CA 90095, USA; Museum of Vertebrate Zoology, University of California, Berkeley, CA 94720, USA; Department of Integrative Biology, University of California, Valley Life Sciences Building, Berkeley; California Academy of Sciences, San Francisco, CA 94118, USA; Center for Tropical Research, Institute of the Environment and Sustainability, University of California, Los Angeles, CA 90095, USA

**Keywords:** annual/perennial, aridity, California, desert plants, diversification, Linanthus, molecular phylogeny, night-blooming, polymorphism, RADseq, target capture

## Abstract

**Premise:** Understanding how arid-adapted plants have diversified in harsh environments is a central question in evolutionary biology. *Linanthus* (Polemoniaceae) occurs in biodiverse dry areas of Western North America and exhibits extensive floral trait variation, multiple color polymorphisms, differences in blooming time, and variation in life history strategies. Here, we reconstruct the evolutionary history of this group.

**Methods:** We generated restriction-site associated (ddRAD) sequences for 180 individuals and target capture (TC) sequences for 63 individuals, with complete species sampling. Using maximum likelihood and pseudo-coalescent approaches, we inferred phylogenies of *Linanthus* and used these phylogenies to model the evolution of phenotypic traits and investigate the geographic speciation history of this genus.

**Key results:** Shallow relationships are consistent and well supported with both ddRAD and TC data. Most species are monophyletic despite rampant local sympatry and range overlap, suggesting strong isolating barriers. The non-monophyly of some species is possibly due to rapid speciation or issues with current species delimitation. Perenniality likely evolved from annuality, a rare shift in angiosperms. Night blooming evolved three times independently. Flower color polymorphism is an evolutionarily labile trait and is likely ancestral. No single geographic mode of speciation characterizes the radiation but most species overlap in range, suggesting they evolved in parapatry.

**Conclusions:** Our results illustrate the complexity of phylogenetic inference for recent radiations, even with multiple sources of genomic data and extensive sampling. This analysis provides a foundation to understand aridity adaptations, such as evolution of flower color polymorphisms, night blooming, and perenniality, as well as speciation mechanisms.

## INTRODUCTION

Desert biomes make up 17% of Earth’s land surface (Millennium Ecosystem Assessment 2005), and appear lifeless for most of the year when lack of precipitation and high temperatures hamper plant survival and growth. However, the desert comes alive in an explosion of color after it rains, when dormant seeds germinate and flowers form thick brushstrokes of pigment. This uncommon and brief outburst of life is made possible by the annual angiosperms, which await suitable environmental cues to germinate in mass. In California, deserts make up 38% of the area (Mooney and Zavaleta 2016), with annuals making up a large part of the species diversity at 52% of total plant species in these deserts (Calflora 2023). This overrepresentation of annuals in the desert and their varied adaptations make them ideal organisms to study the patterns and processes of diversification in this harsh environment.

The deserts of California and their diversity of annual species seem to be young on a geologic time scale (Thorne 1986). Recent meta-analyses of California plant diversity support this idea. For instance, Kraft et al. (2010) showed that the California desert regions have a high concentration of young species with restricted geographic ranges, while Thornhill et al. (2017) found a significant concentration of short phylogenetic branches (i.e., recent diversification events) restricted to the California deserts. However, how deserts facilitate the rapid diversification of plant species remains poorly understood. Habitat heterogeneity, large ranges with isolated populations, and a broad diversity of adaptations to xeric landscapes have all been proposed as potential drivers for high rates of speciation in desert species (Stebbins 1952). The evolution of the annual habit seems to be primed by unstable environments with dry conditions and unpredictable rainfall (Friedman 2020), which is typical in deserts. Annual plants have a fast rate of evolution which is correlated with their short life cycle, isolated populations, and variable environment (Smith and Donoghue 2008; Smith and Beaulieu 2009). This suggests that deserts may promote the evolution of new annual plant species (Stebbins 1952). This is relevant because the correlation between harsh environments and plant diversification has been understudied (Stebbins 1952, but see Hernández-Hernández et al. 2014; Singhal et al. 2021; Lichter-Marck and Baldwin 2022).

There are few phylogenetic studies of California desert annuals. Most published studies do not include all members of the focal clades and almost never include multiple individuals per species, limiting understanding of the patterns and processes of intra and interspecific diversification. Additionally, most studies have only used a handful of loci, largely resulting in poorly resolved phylogenies (e.g., Spencer and Porter 1997; Moore and Jansen 2006; Evans et al. 2009; Porter et al. 2010; Cacho et al. 2014; Walden et al. 2014; Azani et al. 2019; Vasile et al. 2020). More recently, a limited number of studies have used genomic approaches and more extensive taxon sampling shedding light on species-level relationships, patterns of within-species genetic variation, and potential drivers of diversification (Simpson et al. 2017; Mabry and Simpson 2018; Lichter-Marck et al. 2020; Pearman et al. 2021; Rose et al. 2021; Singhal et al. 2021). Given that deserts are young hotspots of biodiversity, well-sampled groups are needed to assess the monophyly of lineages, elucidate species relationships, and untangle complex evolutionary patterns, typical of recent radiations. This is especially worthwhile in geographic regions where species overlap extensively in their geographic range with ample opportunities for gene flow. Such robust phylogenetic studies could inform our understanding of the evolution of traits, adaptations to extreme environments, and whether aridity can be a stimulus to evolution (Stebbins 1952).

The genus *Linanthus* Benth. (Polemoniaceae) is an ideal system to study species diversification in the heterogeneous arid environments of southwestern North America. Half of the currently recognized species co-occur in a geologic transition zone characterized by exceptional plant endemism and environmental variation in Southern California (Kraft et al. 2010). Nineteen out of 25 species overlap in geographic range, and at least fourteen species pairs co-flower and co-occur at a local scale (pers. obs.). Reproducing in a span of a few weeks in the spring, sympatric *Linanthus* species likely have strong reproductive isolation mechanisms to maintain their genetic and phenotypic integrity. *Linanthus* species also display extensive interspecific diversity in habit, blooming time, flower color, and floral scent. These traits attract a diverse suite of pollinators, including beetles, moths, butterflies, hoverflies, long tongue flies, and bees (Chess et al. 2008; Rose and Sytsma 2021; pers. obs.), which likely facilitate the reproductive differentiation of species (Fig. 1). In addition, seven *Linanthus* species have extremely restricted geographic ranges and are ranked as rare, threatened, or endangered in California, including *L. bellus, L. bernardinus, L. concinnus, L. jaegeri, L. killipii, L. maculatus,* and *L. orcuttii* (California Native Plant Society). The extensive sympatry of species, their diversity of pollinator attraction strategies, and their asymmetric geographic ranges point to a complex speciation history that may include speciation with gene flow, micro-allopatry, parapatric speciation, and ecological isolation. However, these phenomena have not been examined in detail.

**Figure 1.**
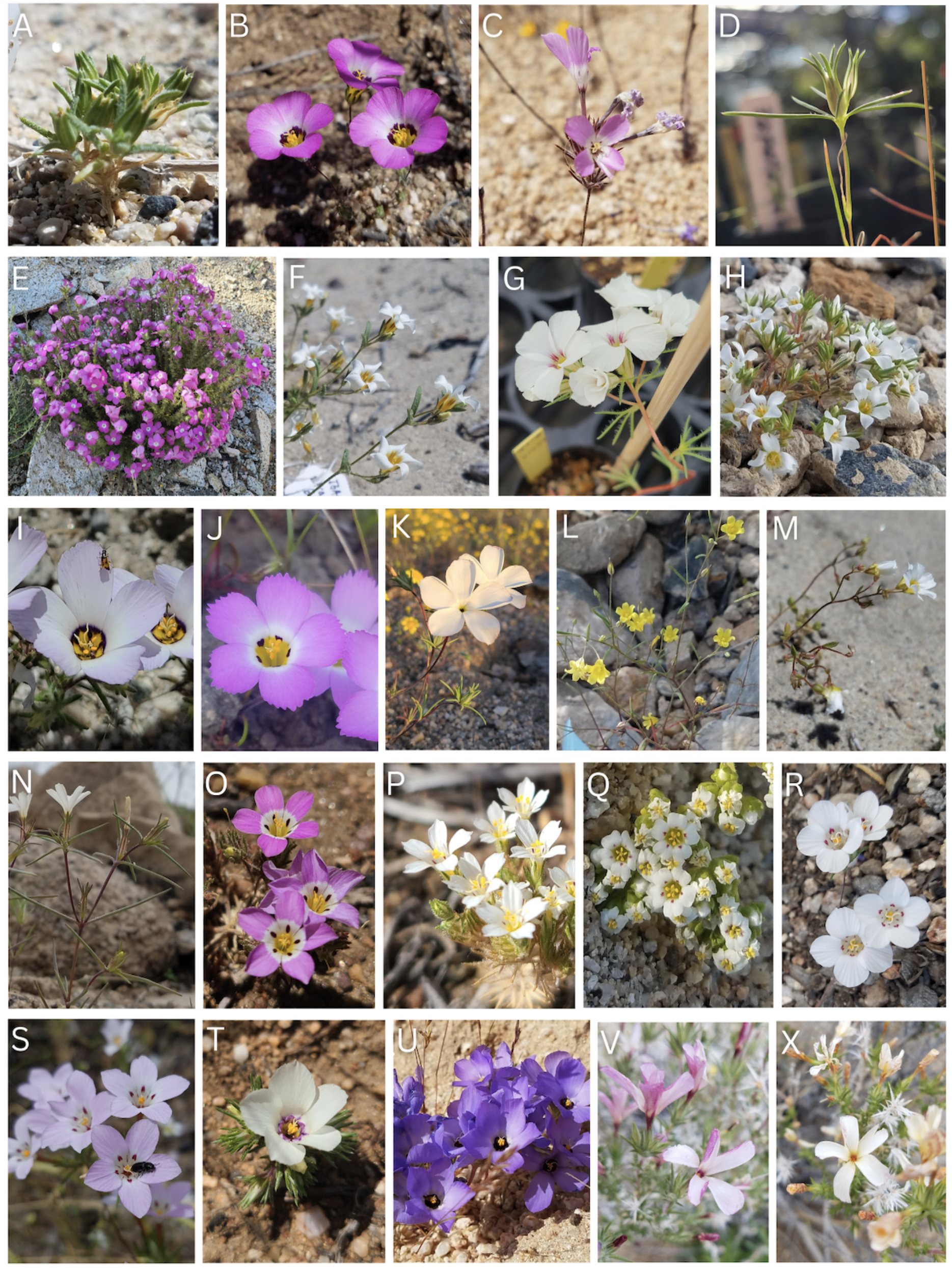
*Linanthus* encompasses extensive floral diversity with many polymorphic species in color and floral markings. Species greatly differ in corolla tube depth and in pollinators observed to visit flowers. Photographs were taken by I.G.A. (A) *L. arenicola.* (B) *L. bellus.* (C) *L. bernardinus.* (D) *L. bigelovii.* (E) *L. californicus.* (F) *L. campanulatus.* (G) *L. concinnus.* (H) *L. demissus.* (I,J) *L. dianthiflorus.* (K) *L. dichotomus.* (L) *L. filiformis.* (M) *L. inyoensis.* (N) *L. jonesii.* (O,P) *L. killipii.* (Q) *L. maculatus.* (R,S) *L. orcuttii.* (T,U) *L. parryae.* (V,X) *L. pungens*.

*Linanthus* species also show extensive intraspecific variability in color and nectary markings, with individual plants taking on different phenotypes in the same population or in disparate areas of the species range. Eleven of the 25 species are color polymorphic, with both white and colored individuals across their range (Fig. 1I&J, O&P, R&S, T&U, V&X). Seven of these color-polymorphic species exhibit within-population color variation, with both colors present in the same population (pers. obs.). One notable example is *Linanthus parryae,* the purple and white polymorphic species at the center of the classic evolutionary debate investigating whether natural selection or genetic drift maintains intraspecific polymorphisms. Different color morphs seem to fare better in wet or dry years, a potential adaptation to the variable desert environment (Epling and Dobzhansky 1942; Wright 1943; Schemske and Bierzychudek 2007). Another potential strategy of several *Linanthus* species to survive in arid environments is night anthesis, where flowers are only open at night and closed during the day during the flowering season. It is not known whether these desert adaptations have facilitated the diversification of *Linanthus*.

*Linanthus,* as currently recognized, includes 25 species (Porter and Johnson 2000) that have diversified over the last 5 Ma and is sister to a clade that includes the genera *Leptosiphon* Benth. and *Phlox* L. (Bell and Patterson 2000). Previous phylogenetic work for 15 species using the nuclear ribosomal internal transcribed spacer (ITS) recovered low support for most relationships across the group (Bell et al. 1999). Another ITS and matK phylogeny that sampled 17 species did not provide a better resolution into the phylogenetic relationships between species (Bell and Patterson 2000). A recent study using 14 nuclear loci for 22 species (with two samples per species) inferred a well-resolved phylogeny and found that eight species were not monophyletic (Landis 2016). However, this study lacked complete taxonomic and broad geographic sampling per species. A robust phylogeny is needed to provide a backbone for tackling inter and intraspecies evolutionary questions in this group.

In this study, we present the first *Linanthus* phylogeny with complete species sampling and broad intraspecific sampling, using two types of genomic data, double-digest restriction-site associated sequencing (ddRADseq) (Peterson et al. 2012) and target capture (TC) of 353 nuclear angiosperm specific genes (Johnson et al. 2019). Our ddRAD data further include multiple samples from 17 species that co-occur locally with a congener to examine the potential for gene flow between sympatric individuals from different species. We use the resulting phylogenies to examine the monophyly of species, reconstruct evolutionary relationships, and investigate patterns of floral evolution and life history shifts. We also use the ddRAD data to study the population structure within selected clades to better understand existing species delimitations for several species. Lastly, we explore the geography of speciation across *Linanthus* to determine the most likely speciation mode in this remarkable desert radiation.

## MATERIALS AND METHODS

### Data collection and processing—

We included representative samples from all 25 currently recognized species (Moran 1977; Porter and Johnson 2000; Porter and Patterson 2015) and outgroups from the genera *Leptosiphon* and *Phlox* (Landis et al. 2018). We collected 83 individuals in the field across California and Nevada and sampled 125 individuals from eight herbaria across the western United States (Appendix S1, see Supplemental Data with this article). We stored the field-collected tissue in silica gel until storage at –20C or in liquid nitrogen until storage at –80C in the laboratoty at UCLA. Some *Linanthus* species are minute; therefore, we carefully dissected all collections to ensure we selected tissue from only one individual. In total, we included samples from 50 individuals that co-occurred with a congeneric species, either observed by us in the field or when herbarium specimens came from the exact same location. We included sympatric individuals for most sympatric species pairs. Seventeen of the species sampled co-occurred with a congener, with a total of 24 combinations of species pairs. See Appendix S1 for complete sample metadata.

We collected genomic data using two approaches, double-digest restriction-site associated sequencing (ddRADseq) and target capture (TC). RAD sequencing generates data that has been used to successfully address questions in both population genomics and phylogenetic relationships in closely related species (Rubin et al. 2012; Eaton et al. 2016, Jacobs et al. 2021). We used a double-digest RAD approach which uses two-enzymes to better recover the same fragments of DNA across samples and reduce sequencing costs (Peterson et al. 2012). In total, we sampled 192 individuals across the 25 currently recognized species and 4 outgroups in *Phlox* and *Leptosiphon*, at an average of 7.2 individuals sampled per species, to represent the species’ breadth of geographic range and phenotypic diversity. The number of individuals sampled per species ranged from 1 to 17, proportional to the species geographic range size (Appendix S2, S3).

For TC, we use the Angiosperm353 bait set (Johnson et al. 2019). This approach can sequence both exons and their flanking regions providing informative phylogenetic data at various phylogenetic scales (Larridon et al. 2020; Slimp et al. 2021). We sampled 63 individuals across 24 currently recognized species and 4 *Leptosiphon* outgroups. We dropped one species (*L. uncialis*) because DNA extraction did not generate enough DNA. We sampled a range of 1 to 5 individuals per species, with an average of 2.5 individuals sampled per species of *Linanthus*.

### DNA extraction, library preparation and sequencing—

We extracted genomic DNA using a modified CTAB technique (Doyle and Doyle 1987; Cullings 1992) that includes an additional incubation step to remove pectin. We prepared sequencing libraries at the Evolutionary Genomics Laboratory of the Museum of Vertebrate Zoology at UC Berkeley.

*ddRAD*—For the RAD-seq libraries, we followed a modified version the 3RAD protocol (Bayona-Vásquez et al. 2019) with adapters and indexing oligos provided by the Glenn lab at the University of Georgia (https://baddna.uga.edu/). We normalized a total of 192 samples to 125 ng of DNA in 10 µL volume and used this as input for a combined digestion and ligation reaction. We used XbaI andEcoRI-HF restriction enzymes to digest the genomic DNA, and the third enzyme (NheI-HF) served to digest adapter dimer produced during the ligation stages of the reaction. Adapter-ligated samples were purified using Solid-phase reversible immobilization (SPRI) beads (Rohland and Reich 2012; Jolivet and Foley 2015). We then amplified the adapter-ligated libraries with indexing oligos (Glenn lab, University of Georgia) and the KAPA HiFi PCR Kit (Roche, Indianapolis, Indiana, USA) using 16 cycles of PCR, following this with another SPRI bead cleaning. We quantified the libraries, pooled them in equimolar amounts, and then size selected fragments at a length of 375-525 base pairs (bp) using a Pippin Prep (Sage Science, Beverly, Massachusetts, USA) at the Functional Genomics Laboratory at UC Berkeley. We quantified the final size-selected library pool with the Qubit® Fluorometer with dsDNA High Sensitivity Assay Kit (Thermo Fisher Scientific, Waltham, Massachusetts, USA) and checked for quality using a Bioanalyzer DNA1000 kit (Agilent Technologies, Santa Clara, California, USA). We sequenced libraries on one lane of Illumina NovaSeq SP 150PE at Vincent J. Coates Genomics Sequencing Lab (GSL) at UC Berkeley (Berkeley, California, USA) for a total of 115 Gb of data at 20x coverage.

*Target capture*—We fragmented DNA from 63 samples, ranging from 175–1100 ng to a target size of 350 bp using a qSonica sonicator (Newton, Connecticut, USA). We sonicated samples with high-quality DNA for 9 minutes at 40% amplitude with a 15s on/15s off pulse. Samples that had a variety of fragment sizes were sonicated for three minutes, while those that were already highly fragmented were not sonicated at all. We cleaned the sonicated DNA and size selected with a double-sided SPRIbead cleaning using 0.525x for right-side selection and 0.675x for left-side selection. We prepared uniquely dual-indexed libraries for Illumina sequencing using the KAPA HyperPrep Kit (Roche, Indianapolis, Indiana, USA) using one-quarter reagent volumes, with a custom-designed dual indexing oligo set developed at the Functional Genomics Laboratory at UC Berkeley. For capture hybridization we enriched 4 µg pools containing equal mass of 7-9 samples each at 62°C for forty hours using the Angiosperm 353 gene set (Johnson et al. 2019) ordered from Arbor Biosciences (“Angiosperms 353 v1”, Catalog #308108.v5) and their myBaits Target Capture v.5 reagents and protocol (http://www.arborbiosci.com/mybaits-manual, Arbor Bioscience, Ann Arbor, Michigan, USA). We amplified enriched products with KAPA HiFi 2X HotStart ReadyMix PCR Kit for 10-12 cycles. We checked the resulting libraries for quality with an Agilent Bioanalyzer DNA1000 assay (Agilent Technologies, Santa Clara, California, USA) and quantified them with the Qubit Fluorometer with dsDNA High Sensitivity Assay Kit. We sequenced 150 paired-end reads at QB3 Genomics on a partial lane of an Illumina NovaSeq S4 for a total of 80 Gb of data.

### Assembly—

*ddRAD*— The sequencing facility demultiplexed the raw sequence reads and quality-checked them with FASTQC v0.11.9 (http://www.bioinformatics.babraham.ac.uk/projects/fastqc/). We used ipyrad to assemble the ddRAD reads using a reference genome for *Linanthus parryae* (Eaton and Overcast 2020; Anghel et al. 2022). We trimmed the variable length adapter sequences using cutadapt (Martin 2011). We filtered low quality base calls with a phred Qscore of 33 and allowed up to five low quality bases per read. We set the minimum length of the reads after trimming to 35 bp, and a stricter filter for adapters at two. We set the clustering threshold within samples to 94% and between samples to 90%. We set the minimum number of samples per locus to four (see full parameters in Appendix S4). After filtering, we used 180 samples in the final assembly.

*Target capture*— The sequencing facility demultiplexed the reads, and we trimmed adapter sequences and quality filtered them using Trimmomatic (Bolger et al. 2014). We assembled sequences for using HybPiper (Johnson et al. 2016) with the mega353 target file (McLay et al. 2021). We assembled flanking regions using ‘intronerate.py’ in HybPiper and identified paralogs using the ‘paralog_investigator.py’ script. The output of HybPiper was exons, supercontigs, paralogs and introns. We removed sequences found to be paralogous and used supercontigs for further analysis because they contain both exons and flanking regions, which can provide information on both the conserved gene regions and the more variable non-coding regions. We used MAFFT to align supercontigs (Katoh et al. 2009). We assessed the occupancy of samples in supercontig alignments with ‘phyluce_align_get_only_loci_with_min_taxa’ function in PHYLUCE (Faircloth 2016) and kept only supercontigs found in all samples for further analyses.

### Phylogenetic inference and species trees—

*ddRAD*—We inferred phylogenetic trees using an alignment of concatenated loci. We used a maximum likelihood analysis approach in IQ-TREE with the GTR model accounting for ascertainment bias (Minh et al. 2020). The ascertainment bias model is used for types of data that do not contain constant sites, such as single-nucleotide polymorphisms (SNPs), to prevent overestimation of branch lengths (Lewis 2001). We assessed support with 1000 ultrafast bootstrap replicates. We reconstructed phylogenetic trees with a minimum of 4, 9, 18, and 36 samples per locus to assess the role of missing data in recovering topologies. Within– and between-species relationships were consistent between these data sets. Results of analyses with the data set using a minimum of 4 samples per locus are shown here. We inferred species trees with SVDQuartets (Chifman and Kubatko 2014) in PAUP* (Swofford 2002). To obtain branch lengths under the multispecies coalescent, we used the qAge command (Peng et al. 2022). As for the concatenated analyses, we estimated species trees with matrices including a minimum of 4, 9, 18, and 36 samples per locus. The species tree recovered with the matrix using a minimum of 4 samples per locus is presented in the results.

We also reassembled the data to include one sample per species to generate a species tree to use for downstream analyses where only one tip represents the species. To do this, we chose the sample within a species with the highest number of recovered loci. If the species was not monophyletic (see Results), we chose the sample with the highest number of loci that belonged to the clade including most samples for such species. We built this tree in IQ-TREE with the GTR model accounting for ascertainment bias and assessed support with 1000 ultrafast bootstrap replicates.

*Target capture*—We inferred gene trees using the most conservative data set of 219 supercontigs with 100% occupancy using IQ-TREE with 1000 bootstrap replicates and the GTR + F default model of substitution (Minh et al. 2020). Then, we used newick utilities to collapse poorly supported gene tree nodes (ML bootstrap support < 0.2) into polytomies (Junier and Zdobnov 2010). We used the gene trees as input for ASTRAL-III v5.7.8 (Zhang et al. 2018) to infer a species tree for all 63 taxa as well as for the 24 species by assigning taxa to species a priori. We also estimated a phylogenetic tree with a concatenated matrix of the loci with 100% occupancy for the 63 taxa in IQ-TREE.

### Population structure—

To assess population structure within certain clades where the taxonomic assignment pointed to non-monophyletic species (see Results), we used rmaverick v1.0.5 (Verity and Nichols 2016). We used only the RAD data for these analyses, with an average of seven samples per species. To prepare files for input into rmaverick, we processed the ipryrad Structure file following methods outlined at https://github.com/zapata-lab/ms_rhizophora/blob/main/analysis_code (Aburto-Oropeza et al. 2021). We processed the vcf file output from ipyrad using vcftools (Danecek et al. 2011). We removed non-biallelic sites and filtered genotypes called below a certain threshold across all individuals (--max-missing 0.25, 0.50, 0.75) for all the population structure analyses. We did this to compare the effect of missing data on the genetic structure output. We also kept only the center SNP from each locus to avoid effects of linked loci.

In rmaverick, we ran the MCMC sampling every 100 steps, with 10% burn-in, 1000 sampling iterations, 20 rungs, and chose a K range from 1 to n + 1 where n is the number of taxonomic assignments in the focal clade. We ran these analyses for 0.25, 0.50, 0.75 missing data sets, and with and without the admixture model. Comparing the data sets using 0.25, 0.50, 0.75 missing data, we found little difference in the structure cluster assignments or values of K, so we present the 0.50 missing data sets here. Because rmaverick is more accurate than other population clustering programs at estimating the number of subpopulations, we chose to report only the value of K with the highest evidence shown by the largest posterior probability (Verity and Nichols 2016). In all cases, the model without admixture was a better fit to the data.

### Ancestral state reconstruction —

To model the evolution of phenotypic traits, we coded all species for annuality/perenniality, day/night blooming, and lack/presence of corolla lobe anthocyanin pigment polymorphism (Appendix S1). We gathered this phenotypic data using the Jepson eflora, the Flora of North America, the monograph of *Linanthus,* and personal observations (Patterson and Porter 2021; Patterson and Porter in prep.; Danforth 1945). While corolla lobe polymorphic species have white, yellow, pink, lavender, purple, and/or peach forms, we focused only on the anthocyanin pigments. Therefore, we coded species with white, yellow or peach corolla lobes as lacking anthocyanin pigments, and the species with pink, lavender, and purple corolla lobes as having anthocyanin pigments (Tanaka et al. 2008). All species with anthocyanin pigments were polymorphic, with populations or individuals within populations with white corollas.

Using the phylogeny built with ddRAD data and one sample per species, we then fitted a Mk model for discrete character evolution (Lewis 2001). To do this, we used the R package *phytools* v1.9-23 and the functions ‘fitMk’(Revell 2012). We took an agnostic approach in choosing the model of evolution for the perenniality and night blooming traits. For these two traits, using the AIC, we selected the best model of evolution between equal rates, all rates different, and irreversible rates where loss of a trait is possible while its gain is not (Appendix S5). We found that the irreversible model had the highest AIC scores for perenniality and night blooming (Appendix S5). For anthocyanin absence or polymorphism, we chose the all rates different model because of the likely different evolutionary rates between loss and gain of flower pigmentation (Rausher 2008), though we report stochastic character density maps with alternate models of evolution (Appendix S6). We simulated stochastic character maps onto the phylogenetic tree and a hundred simulations (Huelsenbeck et al. 2003). Using the ‘densityMap’ function, we visualized the stochastic mapping posterior probability density as a color gradient along the branches of the tree. Then, using the ‘density’ function, we calculated the relative distribution of state changes from the stochastic mapping across the tree.

### Geography of speciation —

To explore the geographic speciation history of *Linanthus*, we used data on species geographic ranges and phylogenetic divergences (Barraclough and Vogler 2000). Specifically, we estimated the relationship between phylogenetic distance and geographic range overlap between all species pairs to evaluate evidence for a predominant geographic (allopatric vs. sympatric) mode of speciation in *Linanthus*. Under this approach, if allopatric speciation is the dominant process, geographic range overlap between young species pairs should increase from ca. 0% to random association as species pairs become more divergent over time. By contrast, if sympatric speciation is the dominant process, geographic range overlap should be ca. 100% between young species pairs but decrease over time among older pairs due to post-speciation geographic range changes (Losos and Glor 2003; Fitzpatrick and Turelli 2006; Skeels and Cardillo 2019). To determine species geographic ranges, we downloaded species location data from the Southwestern Environmental Information Network (SEINet 2022) and the California Consortium of Herbaria (CCH 2022), using only records backed by herbarium collections. We filtered the data to exclude records without precise GPS coordinates (fewer than two decimals precision) and clear outliers in terms of known species distributions. We used these records to create range maps for each species and estimate geographic range overlaps between species pairs using the R package *hypervolume* (Blonder et al. 2014) and the approach implemented at https://github.com/eliotmiller/ebirdr/blob/master/R/hypervolumeOverlaps.R (Miller et al. 2019). Because hypervolumes are geometric shapes with complex geometries, including the presence of holes (Blonder 2016), they can describe species geographic ranges more faithfully beyond simple ellipsoids or convex hulls. Areas of the species range perimeter with no occurrence points can be excluded and ranges can have disjunctions (Blonder et al. 2014).

Consequently, this method enables estimates of range overlap in heterogeneous environments with potential for small-scale allopatry. We calculated the Sørenson similarity index for each species pair and recorded the values in a similarity matrix with values of 0 representing no range overlap and values of 1 representing complete overlap. We used the phylogeny built with ddRAD data and one sample per species to calculate the phylogenetic distance between all species pairs using the ‘cophenetic’ function in the R package *ape* (Paradis et al. 2004). We coded species pairs belonging to the same clade as “within clade comparisons” and to different clades as “between clade comparisons”. We then fitted linear regressions with the ‘lm’ function in R across all species pairs as well as within and between clade comparisons.

## RESULTS

### Data processing—

The ddRAD sequencing yielded an average of 2,440,318 reads per sample. The assembly with 180 samples in a minimum of four individuals yielded 36,861,279 bp, 3,131,821 SNPs, and a total of 165,943 loci with 95% missing data. An average of 7,646 loci per sample were retained. The assembly using one sample per species yielded 4,047,394 bp, 459,834 SNPs, 17,751 loci in a minimum of four individuals, and an average of 4,053 loci per sample. For the TC data, we selected a total of 219 contigs with 100% taxon occupancy for downstream analyses.

### Phylogenetic inference—

The ddRAD and the TC data sets both resulted in well-resolved phylogenies. In the concatenated ddRAD maximum likelihood phylogeny, all nodes had a bootstrap value above 95, with only two nodes at 73 and 76 (Fig. 2A). With an average of seven samples per species, the majority of species resolved as monophyletic (Fig. 2B). The TC phylogeny inferred using IQtree (Fig. 3A) recovered nearly all of the same species relationships as in the phylogeny using ddRAD (Fig. 2A). There were two exceptions in the congruence of the ddRAD and TC phylogenies. One is in a clade recovered with ddRAD data that included *L. bellus, L. concinnus, L. dianthiflorus, L. orcuttii,* and *L. uncialis*. The TC data resolves *L. dianthiflorus* and *L. concinnus*, and *L. bellus* and *L. orcuttii* as a grade, but no samples of *L. uncialis* were included in the TC analysis. The other incongruence was in a clade recovered with ddRAD data that included *L. demissus*, *L. bernardinus*, *L. killipii*, and *L. parryae.* Target capture data supports *L. bernardinus*, *L. killipii*, and *L. parryae* as more closely related to the annual night blooming clade than to *L. demissus*. The ddRAD and TC data represent two independent sources of data and some incongruence between the evolutionary history of these two types of genetic data might be expected due to incomplete lineage sorting. Given that our results were generally consistent across data sets, we are informally referring to several common clades and one grade as follows.

**Figure 2.**
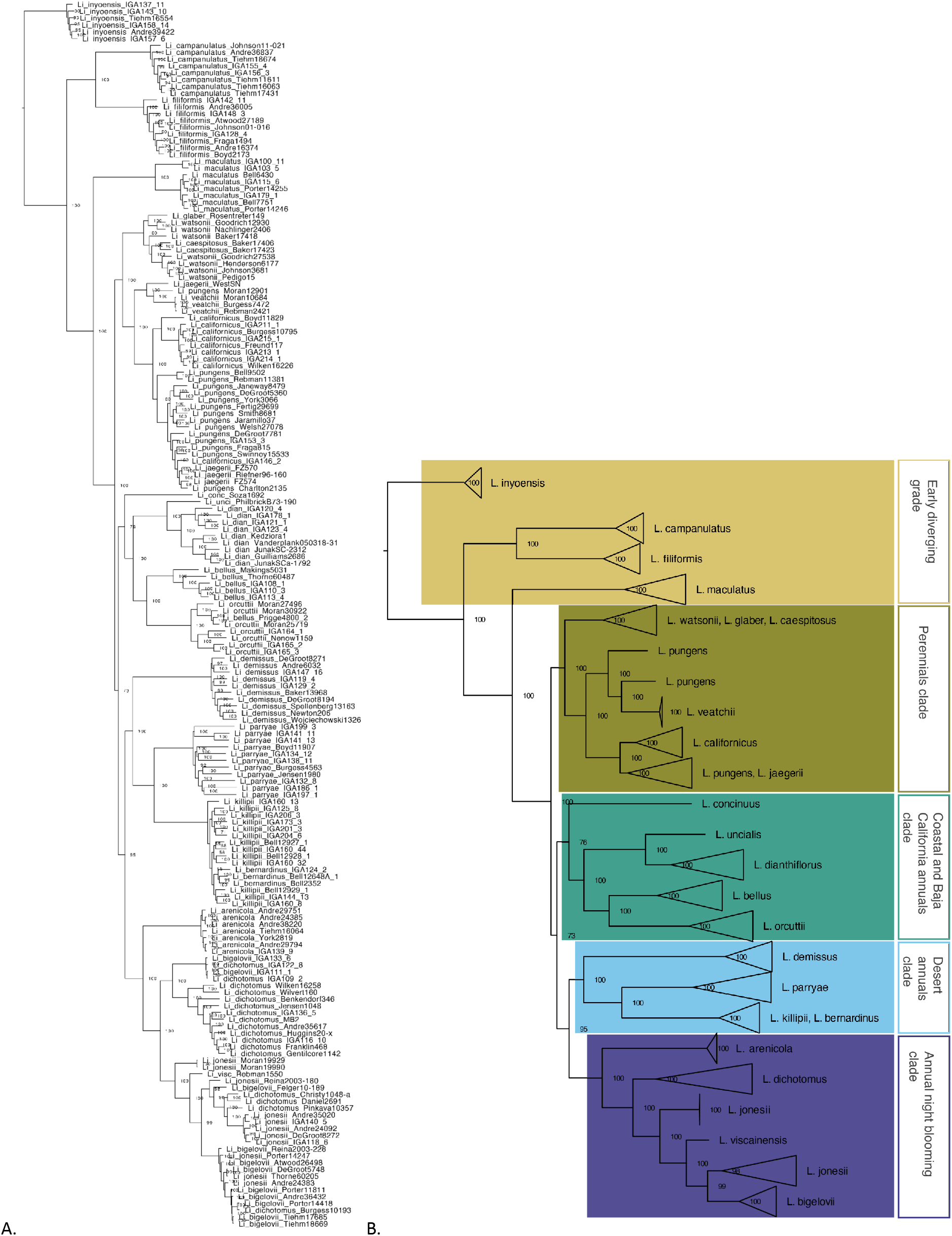
The phylogeny of *Linanthus* inferred using ddRAD data is well resolved and highly supported. A. Maximum likelihood phylogenetic tree inferred in IQtree including 180 samples for all species of *Linanthus* using a concatenated matrix of ddRAD data and a minimum of 4 samples per locus. Values at nodes represent bootstrap support. The average of 7 samples per species included shows that most of the species are monophyletic. B. The same phylogenetic tree as in A with collapsed species. The clades recovered share common morphological features, habitats, or habits.

**Figure 3.**
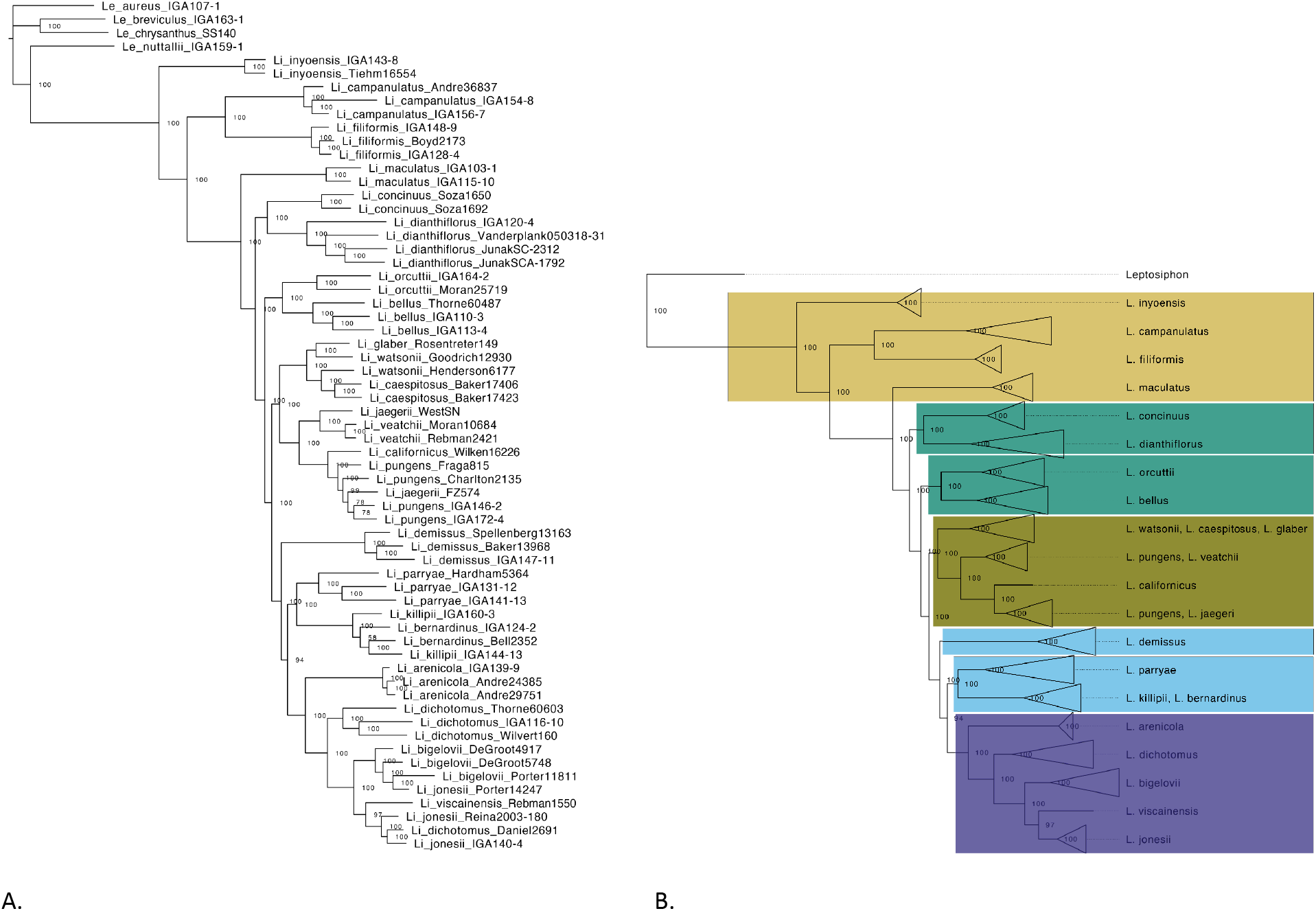
The phylogeny of *Linanthus* inferred using target capture data (Angisoperm 353) data is well resolved and highly supported. This phylogeny is highly consistent with the phylogeny inferred using ddRAD data (see Fig. 2). A. Maximum likelihood phylogenetic tree inferred in IQtree including 63 samples across 22 species of *Linanthus* and 4 outgroups using a matrix of 219 concatenated Target Capture (Angiosperms353) loci with 100% occupancy. Values at nodes represent bootstrap support. Most species are recovered as monophyletic. B. The same phylogenetic tree as in A with collapsed species.

### Early diverging grade—

This grade includes the following species previously described as *Gilia* section Giliastrum: *L. inyoensis, L. campanulatus*, *L. filiformis,* and *L. maculatus* (Grant 1959; Bell et al 1999). While these species did not form a monophyletic group, the grade relationships are well-supported and identical in the ddRAD and TC phylogenies (Figs. 2, 3). The relationships were also consistent with previous studies with limited sampling (Bell et al. 1999; Bell and Patterson 2000). All five species in this grade were recovered as monophyletic (Figs. 2, 3).

### Perennial clade—

Both the ddRAD and TC analyses support the monophyly of a group including the perennial species *L. caespitosus, L. californicus, L. glabrus, L. jaegeri, L. pungens, L. veatchii*, and *L. watsonii* (Figs. 2, 3). These species were previously recognized as the genus *Leptodactylon* Hook. & Arn. (Rydberg 1906). Although a single origin of the perennial clade was also supported in a recent study (Landis 2016), it was inconclusive in earlier phylogenetic analyses (Bell and Patterson 2000). The perennial clade included two nested subclades. One subclade included *L. caespitosus, L. glaber,* and *L. watsonii*, all of which occur outside of California in the Western United States (in Arizona, Colorado, Idaho, Montana, Nebraska, Nevada, Utah, Wyoming). The other subclade included *L. californicus, L. jaegeri, L. pungens*, and *L. veatchii*, all of which occur within California and Baja California, except *L. pungens* which is widespread throughout the Western United States and Mexico.

Most species in the perennial clade were not recovered as monophyletic (Figs. 2, 3). Although *L. caespitosus* and *L. glabrous* were recovered as monophyletic (however, note the small sample size), both species were nested within a paraphyletic *L. watsonii*. In the other subclade, both *L. veatchii* and *L. californicus* were recovered as monophyletic, but they were nested within a clade that included the paraphyletic species *L. pungens* and *L. jaegeri*.

The rmaverick population structure analysis for the perennial clade showed the highest posterior probability for three distinct genetic clusters (Verity and Nichols 2016; Fig. 4). One cluster included all the samples of the species *L. watsonii* and *L. caespitosus*, which occur outside of California, another cluster included all the samples of the Baja California endemic *L. veatchii* and two samples of *L. pungens* from Southern California and Baja California, and a third cluster included all samples of the species *L. pungens, L. californicus,* and *L. jaegeri* (Fig. 4). These results are consistent with the phylogenetic results showing a lack of genetic cohesiveness of the currently recognized taxonomic species in this clade.

**Figure 4.**
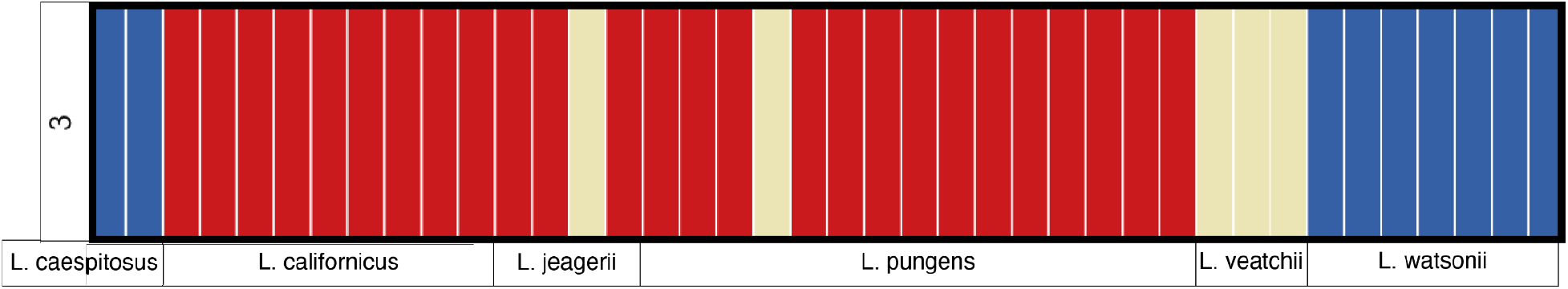
Genetic clusters did not match species taxonomy in the perennial clade. K=3 had the highest posterior probability, meaning that three genetic clusters were recovered for six taxonomic species. The red cluster included all samples identified as *L. californicus* and most samples identified as *L. pungen*s and as *L. jaegeri*. The blue clade included all samples identified as *L. caespitosus* and *L. watsonii*. The tan group included all samples identified as *L. veatchii*, one sample identified as *L. jaegeri*, and one sample identified as *L. pungens*.

### Coastal California and Baja California annuals clade—

The ddRAD phylogeny recovered a monophyletic group including *L. bellus, L. concinnus, L. dianthiflorus, L. orcuttii,* and *L. uncialis*, all of which occur in the coastal or higher elevation areas of Southern California and Baja California (Figs. 2,3). A recent phylogenetic study also recovered this clade (Landis 2016). While the monophyly of this group was not fully supported in the TC phylogeny, all species within this clade were closely related and formed a grade (Fig. 3). The ddRAD analysis with multiple samples per species recovered all species as monophyletic, though we could only include one sample for *L. concinnus* and *L. uncialis* (Fig. 2).

### Desert annuals clade—

In the ddRAD phylogeny, *L. bernardinus, L. demissus, L. killipii,* and *L. parryae* formed a clade sister to the annual night-blooming clade (see below) (Fig. 2). The TC species tree did not support the monophyly of the desert annual clade (Fig. 3). Instead, this tree recovered *L. demissus* as sister to a clade with two subclades, one including the remaining species in the desert annual clade (*L. bernardinus, L. killipii,* and *L. parryae*) and a subclade including the annual night-blooming species (see below) (Fig. 3). The broad sampling in the ddRAD phylogeny showed that most species in the desert annual clade were recovered as monophyletic, with the exception of *L. bernardinus* and *L. killipii*, both of which formed a clade with intermixed samples.

### Annual night blooming clade—

The annual night blooming clade included the species *L. arenicola*, *L. bigelovii, L. dichotomus*, *L. jonesii*, and *L. viscainensis.* All species in this clade are night bloomers, with the exception of *L. dichotomus* where populations of *L. dichotomus* subsp. *meridianus* flower during the day. This group was monophyletic in both the ddRAD and TC phylogenetic analyses (Figs 2, 3). This corroborates the section *Linanthus* described by Grant which included *L. bigelovii, L. dichotomus* and *L. jonesii* (Grant 1959). *Linanthus viscainensis* was later added to the section *Linanthus* based on its morphological similarities to *L. arenicola* (Moran 1977). This clade was also recovered in previous analyses with the addition of one accession of *L. filiformis* (Landis 2016).

Most species in this clade were highly paraphyletic or polyphyletic (Figs. 2, 3). *Linanthus arenicola* was the only monophyletic species. Six out of 11 samples of *L. jonesii* formed a clade, but it was nested within a more inclusive clade with several intermixed samples of *L. dichotomus* and *L. bigelovii*. Eight out of 11 *L. bigelovii* samples formed a clade, but it was nested in a more inclusive clade with several samples of *L. dichotomus* and *L. jonesii*. The only sample of *L. viscainensis* included here was nested within a more inclusive clade including samples of *L. bigelovii, L. dichotomus,* and *L. jonesii*. Thirteen out of 17 *L. dichotomus* samples and two samples of L. bigelovii formed a clade.

The population structure analysis in the annual night blooming clade showed the highest posterior probability for four distinct genetic clusters (Fig. 5). One cluster included the majority of samples identified as *L. bigelovii* (8/11), some samples identified as *L. jonesii* (5/11), and one sample identified as *L. dichotomus* (1/17). A second cluster included the majority of samples identified as *L. jonesii* (6/11), some samples identified as *L. dichotomus* (3/17), and one sample identified as *L. bigelovii* (1/11). A third cluster included only samples identified as *L. dichotomus* (11/17). A fourth cluster included some samples identified as *L. bigelovi* (2/11) and some samples identified as *L. dichotomus* (2/17) (Fig. 5). We did not detect any geographic signal characterizing these genetic clusters.

**Figure 5.**
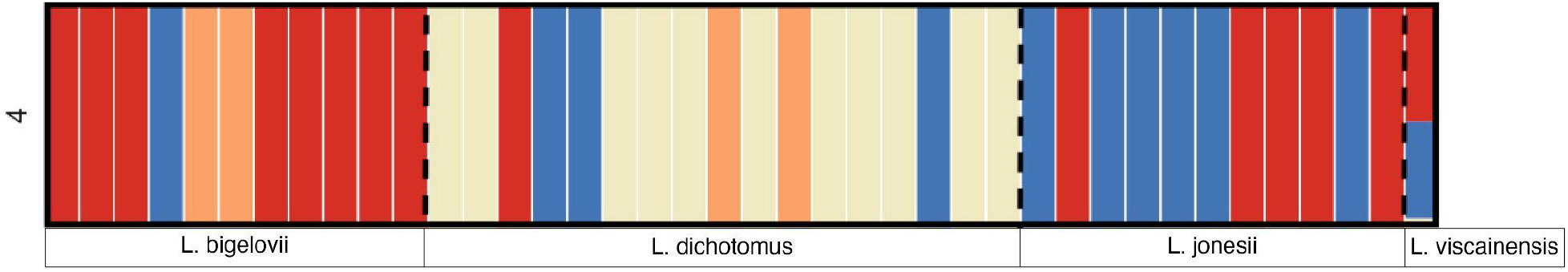
Genetic and taxonomic groups in the annual night bloomers showed little congruence. K=4 had the highest posterior probability, meaning that four genetic clusters were recovered for four taxonomic species. The red cluster included a majority of samples identified as *L. bigelovii*. The blue cluster included a majority of samples identified as *L. jonesii.* The tan cluster included only samples identified as *L. dichotomus*. The orange cluster included two samples identified as *L. bigelovi* and two samples identified as *L. dichotomus*. We excluded *L. arenicola* from this analysis because it was monopholetic.

### Trait evolution and ancestral states —

Mapping the distribution of phenotypic traits onto the ddRAD phylogeny shows that perenniality is clustered in one clade, night blooming appears in the perennial clade and in the annual night blooming clade, and corolla lobe color polymorphisms are dispersed across the phylogeny in three of the major clades (Fig. 6). Stochastic character mapping showed that perenniality evolved once with no reversals to annuality (Fig. 7A). Night blooming evolved three times, once in the annual night blooming clade, and twice in the perennial clade where all but two species have night blooming populations (Fig. 7B, Appendix S7). Corolla lobe anthocyanin pigment polymorphism may be ancestral in *Linanthus*, with several reversals to unpigmented corolla lobes (Fig. 7C). The posterior probability distribution for the number of changes shows that gain of anthocyanin polymorphism may have occurred twice and loss of anthocyanins ten times (Appendix S7).

**Figure 6.**
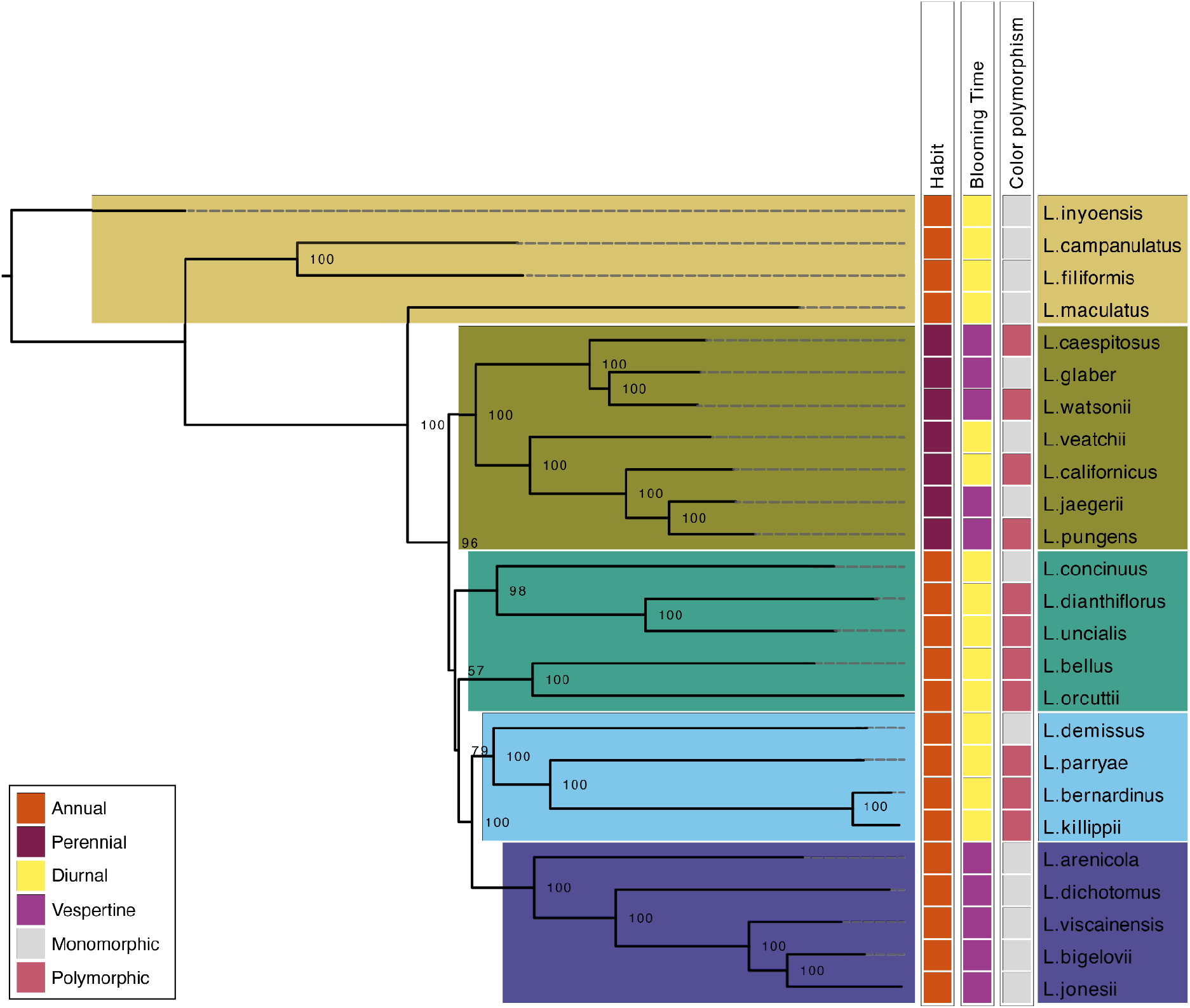
Phylogeny of *Linanthus* showing the distribution of three phenotypic traits. Left, *Linanthus* phylogeny using ddRAD data and one individual per species. Values represent bootstrap support. Right, diagram showing the character states distribution across species. Colors grouping taxa represent the clades we defined in Figure 2B.

**Figure 7.**
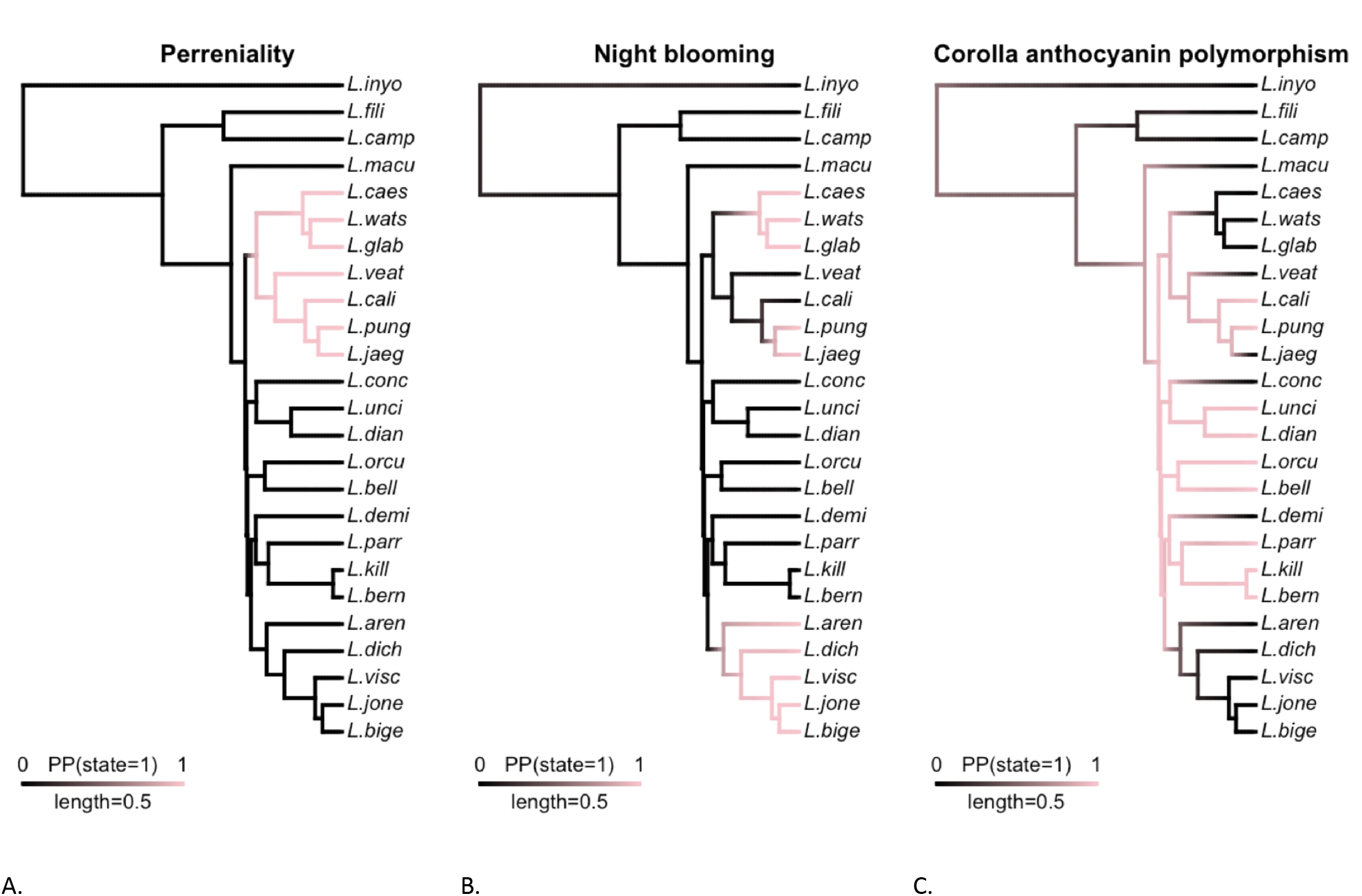
Across the *Linanthus* radiation, some phenotypic traits show little lability while others are highly labile. Each panel shows stochastic character density maps for different traits. A. Perenniality evolved once (1 = perenniality). B. Night blooming evolved three times, once in an annual clade, and twice in the perennial clade (1 = night blooming). C. Corolla lobe anthocyanin pigment polymorphisms may be ancestral in *Linanthus*, with several reversals to unpigmented corolla lobes. The most dominant state across *Linanthus*’ evolutionary history is polymorphic, represented by the dominance of pink color along the branches in the phylogeny.

### Geography of speciation —

The majority of species pairs showed < 50% overlap in geographic ranges regardless of phylogenetic distance (Fig. 8). Nonetheless, species range overlap varied from 0-80 %. Across all species pairs, geographic range overlap decreased with phylogenetic distance, but this relationship was not significant (R^2^ = 0.01533; *P* = 0.1067; Fig. 8). Young species pairs often had non-overlapping geographic ranges and range overlap tended to increase with phylogenetic distance (within clade comparisons). However, this relationship was not significant (R^2^ = 0.00114; *P* = 0.8942; Fig. 8). The relationship between phylogenetic distance and geographic range overlap between older species pairs (i.e., between clade comparisons) was negative and non-significant (R^2^ = 0.003245; *P* = 0.4843; Fig. 8). Together, these results suggest that a single geographic mode of speciation (allopatric or sympatric) does not predominate among *Linanthus* species.

**Figure 8.**
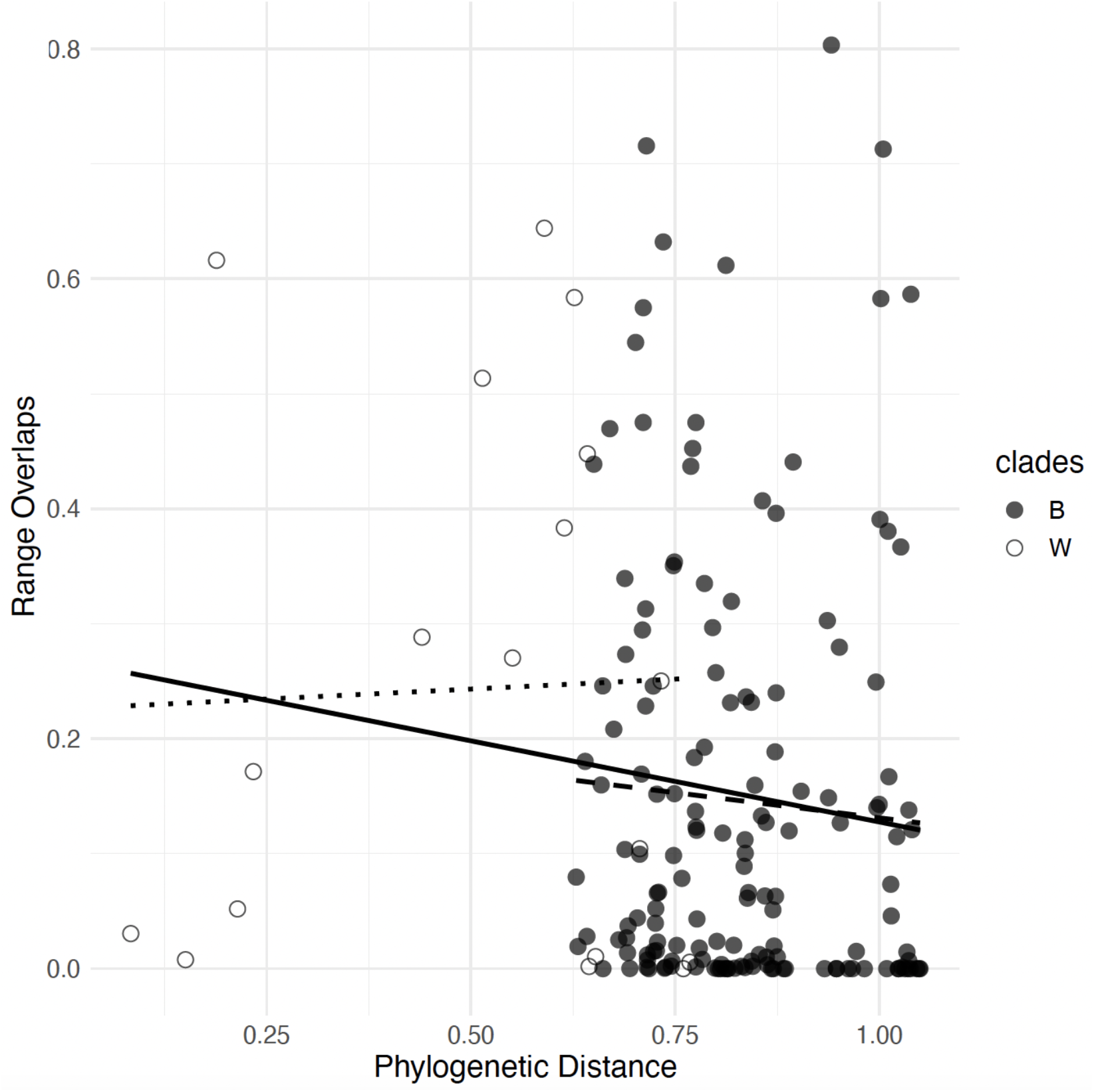
The relationship between phylogenetic distance versus range overlap of pairs of *Linanthus* species shows there is no predominant geographic mode of speciation. Pairwise phylogenetic distance is based on the ddRAD phylogeny using one individual per species, and geographic range overlap is estimated as the overlap in hypervolumes corresponding to geographic ranges for each species. The solid line corresponds to the regression across all pairwise comparisons. The dotted line corresponds to the regression within clade comparisons (W). The dashed line corresponds to the regression between clade comparisons (B).

## DISCUSSION

### Phylogenetic relationships are well-resolved and most species are monophyletic—

Our phylogenetic analyses generated well-resolved phylogenies that included all of the currently recognized species in *Linanthus*. The TC phylogeny (Fig. 3A) recovered nearly all of the same species relationships as in our concatenated ddRAD phylogeny (Fig. 2A), and they both were congruent with results from previous analyses with limited taxon and genetic sampling (Landis 2016). The species tree inferred with SVDQuartets from ddRAD data (Appendix S8) showed inconclusive species relationships due to very short branches, but this method is sensitive to large amounts of missing data common in RAD datasets (Nute et al. 2018). The species tree inferred with ASTRAL from TC data also recovered monophyly of most species, and most shallow species relationships were congruent with the ddRAD data species trees (Appendix S9). Notably, our results showed that most species are monophyletic even when we included multiple samples from different species co-occurring in sympatry with ample opportunities for interspecific gene flow. This suggests that reproductive isolating barriers have likely evolved for most species in this recent desert radiation. The lack of monophyly of some perennial and annual night blooming species may be due to interspecific gene flow, incomplete lineage sorting, or that current species limits are problematic. Further work with deeper taxon and genomic sampling as well as field experiments are needed to rigorously test these hypotheses.

### Perennials evolved from annuals—

The species in the perennial clade all share several synapomorphies including the subshrub habit with a woody base, leaves that grow tightly in fascicles with sharp-tipped and filiform lobes or entire filiform leaves, and salverform corolla. These species occur in mountainous regions, consistent with the finding that alpine environments favor long lived species that can persist the colder winters (Billings and Mooney 1968; Ricklefs and Renner 1994).

Shifts from annual to perennial habit are considered rare across angiosperms, with perenniality thought to be the ancestral state in flowering plants and within most families and genera (Friedman 2020). However, recent phylogenetic comparative work has shown examples of transitions from annual to perennial habit (Tank and Olmstead 2008; Soltis et al. 2013). In Polemoniaceae, transitions between annuality to perenniality in both directions are common including in *Phlox* and *Leptosiphon*, which are two closely related genera of *Linanthus* (Barrett et al. 1996). Both the ddRAD and the TC phylogenies presented here support the evolution of the perennial species from an annual ancestor, and no reversals to annuality (Fig. 7A). Thus, the transition to perenniality is unidirectional from annual ancestors. The evolution of perenniality may have implications for diversification and evolution in *Linanthus*. Perennial plants generally show a slower rate of molecular evolution than annual plants, which has been attributed to longer generation times (Andreasen and Baldwin 2001) and larger population sizes (Bousquet et al. 1992). Diversification rates have also been shown to be lower in perennials than in annuals due to lower extinction rates in annuals (Soltis et al. 2013).

### Night blooming evolved three times—

We found that night blooming has only evolved three times in *Linanthus* (Fig. 7B, Appendix S7) with one reversal to day blooming in *L. dichotomus* subsp. *meridianus*. Night blooming has evolved multiple times across angiosperms but it is relatively rare (Silberbauer-Gottsberger and Gottsberger 1975; Grant 1983). In Polemoniaceae, hawkmoth pollination seems to have evolved in three genera, *Linanthus*, *Ipomopsis* and *Phlox*, but only *Linanthus* has flowers that open exclusively at night (Grant 1983). This suggests further specialization for hawkmoth pollination in *Linanthus* because there are no other pollinators available to visit the flowers at night.

*Night blooming is rare and may be a mechanism of allochronic speciation—*Night blooming species often occur in low densities and are spread across the landscape. This demographic pattern of night bloomers is associated with dry habitats, where daytime anthesis might lead to excessive water loss through transpiration (Stebbins 1970). Populations of the annual night blooming *Linanthus* species show these demographic patterns. If night blooming is indeed advantageous from a hydraulic perspective in *Linanthus*, the question is why more species have not evolved this trait given their widespread distribution in the hot, dry deserts. Detailed physiological and genomic studies could shed light on the biological mechanisms underpinning this phenotype. Alternatively, it is plausible that shifts in flowering time have evolved as prezygotic isolating barriers and are linked to a more complex phenotype involved in pollination and possibly allochronic speciation (Taylor and Friesen 2017). Night blooming flowers are usually white, pale yellow or pale pink with strong scents emitted at night, and with long corolla tubes (Grant 1983; Knudsen and Tollsten 1993). These traits are present in all the annual night blooming *Linanthus*, with variation within and between species, making it a great system to study the genetic basis of this complex phenotype likely involved in speciation. For instance, this system can be used to study which suite of traits is specialized to moth pollination and which attracts a wider variety of pollinators. With their reduction in showy floral features (i.e. lack of pigmentation and petal markings), and fewer pollinators available at night, night bloomers can also be used to investigate whether this strategy is an evolutionary dead end.

Our results show that populations of species in the night blooming clade co-occur in sympatry with populations from species in other clades. Shift in blooming time could work as a temporal isolating barrier when no geographic barriers limit interspecific gene flow. Because night blooming species co-occur with species that have diverged at different times (Figs. 2, 8), it is unclear if night blooming is the cause of reproductive isolation or evolved later in secondary sympatry as a reinforcement mechanism (Taylor and Friesen 2017).

*Selfing may be a mechanism of reproductive isolation between closely related species—*Outside of night anthesis, the *Linanthus* species in the annual night-blooming clade are united morphologically by their white or yellow corolla color, short or non-existing pedicel, and a cylindrical or urn shaped calyx, with the membrane separating sepals that are wider than the lobes. The stamens are included in the corolla tube, with the pistil at or below the stamens. This combination of traits is often found in flowers that can self-pollinate (Ushimaru and Nakata 2002). Indeed, most *Linanthus* species in the annual night blooming clade can set seed without the corolla emerging out of the calyx (pers. obs.). Self-compatibility is common in other hawkmoth pollinated flowers (Grant 1983). For instance, *L. arenicola* and some populations of *L. jonesii* and *L. bigelovii* have very short floral tubes, making hawkmoth pollination less likely.

Therefore, it is possible that some populations of these species are selfers. Further study is needed to determine whether these populations are facultative selfers, growing smaller autogamous flowers in years where conditions are not favorable. The gradient from selfing to outcrossing via hawkmoth pollination in this night blooming clade provides the opportunity to investigate different strategies for reproductive isolation in closely related species. Similar trends in selfing features are present in *Leptosiphon,* a closely related genus of *Linanthus* (Goodwillie 1999; Goodwillie and Stiller 2001; Goodwillie and Ness 2005). Selfing in *Leptosiphon* may have evolved to ensure reproductive success in environments with inconsistent pollinator visitation and with range expansion to drier habitats that decouple pollinator emergence from flowering time (Goodwillie 1997). Whether similar mechanisms operate and evolve independently in *Linanthus* is unknown.

*Scent may act as a reproductive isolating mechanism, especially in night blooming species—* Night blooming species use scent to attract pollinators because visual cues are less effective at night (Raguso and Willis 2002). Many night blooming flowers emit a heavy sweet musky scent, which is an olfactory attractor of noctuid moths (Raguso et al., 2003). *Linanthus dichotomus*, the showiest species in the night blooming clade, emits scents associated with moth attraction, and different chemical profiles between the day and night blooming subspecies (Chess et al. 2008). Analyzing the chemical profiles of *Linanthus* species will improve our understanding of the evolution of the hawkmoth pollination syndrome. Further chemical ecology studies are needed to explore how floral scent works as a potential isolating mechanism in *Linanthus*.

### Color *polymorphism* may be ancestral—

Flower anthocyanin polymorphisms in *Linanthus* may have evolved early in the history of the genus, in the clade that includes most species in the genus except for *L. inyoensis*, *L. filiformis* and *L. campanulatus* (Fig. 7C). Although these three species do not exhibit color polymorphisms, several other genera in Polemoniaceae have color-polymorphic species (Schemske and Bierzychudek 2007). It is possible that color polymorphism or pigmentation evolved in the ancestor of *Linanthus* or even earlier in the history of Polemoniaceae, but those analyses are beyond the scope of this paper. A noteworthy feature of the anthocyanin polymorphisms in *Linanthus* is that no species are pigmented monomorphic (i.e., there are no species with only pink or purple petals), and all pigmented species also have a non-pigmented morph (i.e., a species with pink petals also has individuals with white petals). The pigmented monomorphic trait may be a hidden state in the evolutionary history of *Linanthus* but this is not possible to infer from our analyses. Despite this, the prevalence of polymorphisms in the genus indicates that polymorphisms are a shared derived trait in at least some of the *Linanthus* lineages. It is expected that polymorphisms are rarely retained across speciation events, because the genetic variation responsible for the polymorphism as well as the disruptive selective pressure maintaining the polymorphism must persist through time (Jamie and Meier 2020). While polymorphisms might be a precursor to speciation, polymorphisms that persist through a speciation event must be a result of different forces than those driving speciation (Gray and McKinnon 2007). Small scale habitat differences or temporal fluctuations are possible disruptive forces that do not directly lead to reproductive isolation, and have been credited with maintaining the color polymorphism in *Linanthus* parryae (Schemske and Bierzychudek 2001, 2007). The prevalence of color polymorphisms across the *Linanthus* radiation and the potential for its mechanism to be maintained through speciation events makes this genus an ideal subject to investigate the dynamics between speciation and polymorphisms.

Flower pigmentation evolutionary transitions are common across the angiosperm phylogeny (Rausher 2008; Smith and Goldberg 2015), but the macroevolution of color polymorphisms remains understudied. Flower color polymorphisms are thought to be rare across angiosperms but are common in certain clades like *Antirrhineae* (Lamiaceae) and *Protea* (Proteacea) (Carlson and Holsinger 2015; Ellis and Field 2016). In California occurring Polemoniaceae, 36% of the species exhibit flower color polymorphism (Schemske and Bierzychudek 2007), in *Ipomopsis, Leptosiphon, Linanthus* and *Phlox*. In *Linanthus*, 44% of the species are polymorphic across populations and 28% within populations (pers. obs.). The pink, lavender, and purple color variation in *Linanthus* are likely anthocyanin based pigments (Tanaka et al. 2008). The inability to synthesize anthocyanin pigments has been suggested as a mechanism for producing the white corolla individuals (Warren and Mackenzie 2001), but this mechanism remains to be tested in *Linanthus*. The loss of the color polymorphism seems to have occurred several times in the history of *Linanthus* (Appendix S7), supporting that color is likely more easily lost than gained (Rausher 2008).

Many of the species in *Linanthus* exhibit continuous color variation. For example, the iconic *Linanthus parryae* has been coded as dimorphic in the classic papers investigating the evolution of such polymorphism (Epling and Dobzhansky 1942; Wright 1943; Schemske and Bierzychudek 2007). However, observations in the field across the flowering season indicate that individuals range in intensity of flower color, both within and between populations. In addition, the color of the purple morphs appears to fade with heat or time since anthesis (pers. obs.). These characteristics point to a potentially more complex mechanism for the genetics and plasticity of flower color than previously suspected in *Linanthus parryae*, and in the other *Linanthus* polymorphic species.

While anthocyanin produced pigment influences pollinator attraction, it also has survival functions, including deterring herbivores and responding to abiotic stresses such as UV exposure and drought (Strauss and Whittall 2006). Across angiosperms, flower color polymorphisms are most common in heterogeneous environments, like the Mediterranean or high elevation biomes (Sapir et al. 2021). Experimental studies have shown that in white and pink/purple flower polymorphic species, the pink/purple morphs show greater tolerance to drought and heat stress (Warren and Mackenzie 2001; Coberly and Rausher 2003; Vaidya et al. 2018; Dittmar and Schemske 2023). In *Linanthus parryae* pollinator visitation did not differ between two color morphs, but reproductive success was higher in white flowered individuals in wetter years, and in purple flowered individuals in dry years (Schemske and Bierzychudek 2001). This points to a potential pleiotropic effect of flower color and environmental factors in *L. parryae* (Schemske and Bierzychudek 2007) but can also point to potential linked genes producing these seemingly unrelated phenotypes (Rausher 2008). The white and pink/purple polymorphism commonly found in *Linanthus* may be an important adaptation that allowed plants to tolerate the variable environmental conditions in xeric areas of Western North America and might have been a precursor to its diversification in this geographic area with high UV radiation, drought and heat. The prevalence and the potential ancestral origin of color polymorphisms in *Linanthus* opens promising future avenues for studying the mechanisms that maintain polymorphisms within species and across speciation events.

### No predominant mode of speciation, but range overlap in sister species is indicative of parapatric speciation—

We did not find a relationship between phylogenetic distance and species range overlap, thus we did not detect a prevalent geographic mode of speciation in *Linanthus*. Though current species distributions are not necessarily reflective of the species range at the time of speciation (Losos and Glor 2003), some signature of this pattern is likely present in the current geography of closely related species (Barraclough and Vogler 2000). Allopatric speciation has been considered the dominant mode of speciation (Mayr 1959; Coyne and Orr 2004). If this is the case, we would expect geographic range overlap to increase over time with sympatry arising secondarily after populations have accumulated enough differences and become reproductively isolated. However, we did not detect signals of such a pattern (Fig. 8). Recent studies have suggested that in plants, sympatric speciation is more prevalent than other geographic modes of speciation (Skeels and Cardillo 2019). Under this scenario, we would expect geographic range overlap to decrease due to post-speciation geographic range changes (Losos and Glor 2003).

Yet, our findings for *Linanthus* were not consistent with this pattern (Fig. 8). Taken together, our results indicate that multiple speciation mechanisms are at play in the *Linanthus* radiation and not a single mode predominates. For instance, we found that some young species pairs show considerable geographic range overlap, while others overlap minimally. This suggests that in some cases speciation could have happened in parapatry with opportunities for homogenizing gene flow and in other cases, speciation likely happened in isolation in allopatry. Further geographic sampling at the landscape scale combined with detailed studies in regions of geographic overlap using simulations and population genomic approaches will be essential to confidently discern speciation modes.

Recent studies of California plants suggest that the patterns of geographic range overlap, geographic range asymmetry, and time since divergence between species pairs were not consistent with allopatry as the dominant mode of geographic speciation (Anacker and Strauss 2014; Grossenbacher et al. 2014; Christie and Strauss 2018). However, these studies estimated range overlap using a different approach to the one employed here. We used overlap of hypervolumes that account for density of points and holes in the distribution of species occurrences, and holes in overlap from calculations of sympatry (Blonder et al. 2014). This method is sensitive to overlap at a finer scale, potentially excluding areas of overlap where the potential for gene flow between species pairs is low. Fine scale geographic partitioning or “micro-allopatry” may be common in California native plant species (Anacker and Strauss 2014; Grossenbacher et al. 2014). These previous studies used the overlap between polygons formed by occurrence points, and the difference in range overlap calculation methods could explain the overall discrepancies between our findings and the ones reported for those studies.

The expected pattern of increasing overlap with increasing phylogenetic distance under allopatric speciation may be apparent in a genus that diversified across a more homogenous environment, with fewer barriers to range expansion and contraction. However, in *Linanthus* this pattern may not emerge even if allopatric speciation is common because some species pairs never experience secondary sympatry (Fig. 8). This could result from specialization to certain habitats or pollinators, the inability to expand geographic ranges, or the existence of a heterogeneous environment with barriers to dispersal. These are patterns we commonly see in the ecology and geography of *Linanthus*. Testing these hypotheses will lay the groundwork for future studies exploring speciation mechanisms in *Linanthus* across the harsh North American deserts.

## CONCLUSION

In this study, we present a complete phylogeny of the genus *Linanthus*, a diverse radiation of mostly annual plants from the biodiverse deserts of Southwestern North America. Our phylogeny is the first to date to include complete species sampling and extensive intraspecific sampling. This approach allowed us to explore the monophyly of species and species relationships with increased rigor. Most species resolve as monophyletic despite rampant local sympatry and range overlap, suggesting the presence of strong isolating ecological barriers. The species within the annual night blooming clade and the perennial clade are not monophyletic, and closer taxonomic and population level studies are needed to untangle the evolutionary patterns of those species. Although we do not detect a strong signal for a predominant geographic mode of speciation, most species show some overlap in geographic range regardless of time since divergence. This suggests that some species could have evolved in parapatry, likely in the face of gene flow, while others likely evolved in isolation and never attained secondary sympatry. The strategies that *Linanthus* species have evolved to deal with desert living, including flower color polymorphisms, facultative selfing, night blooming, and annuality, make it a rich system to study how plants can adapt to a drier world.

## Supporting information

Appendix S1_SampleMetadata

Appendix S2_SpeciesTraits

Appendix S3_RADsamples_locations_wrap

Appendix S4_ipyradparams-Li_094090_min750_min4_mindep6_refLiparrCCGP

Appendix S5_StochasticCharacterMapping_AICscores

Appendix S6_Anthocyanins_simmap_postprob_AlternateModels

Appendix S7_Alltraits_simmap_distrib_of_changes

Appendix S8A_RADSpeciesTree_Li_svdq_min4_ipy_bs100_SpeciesTreenoBL

Appendix S8B_RADSpeciesTree_SVDQ_SpTreewBL_Li_min4_bs100_ipy_SpeciesTree

Appendix S9A_TC_ASTRAL_trimmed_supercontings-0p_219genes_PhylTree

Appendix S9B_ASTRAL_trimmed_supercontings-0p_219genes_sptree_clades

## ACKNOWLEDGEMENTS

The authors thank Jade Corpus-Sapida and Sarah Payne for helping with DNA extractions. We are grateful to the curators of ASU, BRY, CAS, RSA, SBBG, SD, UCLA, and UNR herbaria for permission to sample from herbarium specimens, especially to Mare Nazaire, who provided a bulk of the herbarium samples used in this study. We thank Robert Patterson for inspiration and helpful discussions about *Linanthus* taxonomy and evolution, and Mark Porter for helpful *Linanthus* discussions and training on the pectinase DNA extraction protocols. We are grateful to Claudia Henriquez for DNA extraction and library preparation training and support. We thank Sonal Singhal for discussion about alternate analyses when we seemed to hit a dead-end, and Isaac Overcast for help troubleshooting our ipyrad assemblies. We thank Duncan Bell, Maisie Borg, and Steven Serkanic for providing supplemental samples for this project. We are very appreciative to Kathleen Kay for providing comments that strengthened the arguments of this manuscript. Funding for this work was generously provided by the Ecology and Evolutionary Biology Department at UCLA, the UCLA La Kretz Center and Stunt Ranch “Graduate Research Grant”, the Botanical Society of America “Bill Dahl Graduate Student Research Award”, the American Society of Plant Taxonomists “Graduate Student Research Grant”, the California Native Plant Society “Hardman Native Plant Research Award” and Felipe Zapata’s start-up funding. This work is part of Ioana Anghel’s Ph.D. dissertation at UCLA.

## AUTHOR CONTRIBUTIONS

I.G.A and F.Z. designed the study. I.G.A conducted the research, analyzed the data and wrote the manuscript. L.L.S and I.G.A prepared sequencing libraries. I.L.M. assembled the target capture sequence data. All authors contributed to the editing and revision of the manuscript.

## DATA AVAILABILITY STATEMENT

All sequences generated for this work will be available on NCBI Sequence Read Archive (SRA) under the BioProject XXX. Appendices, alignments, trees, character matrix and scripts will be available on <u>github.com/ioanaanghel/Linanthus_phylogeny</u>.

## SUPPORTING INFORMATION

Additional supporting information may be found online in the Supporting Information section at the end of the article.

**Appendix S1**. Sample metadata with location and provenance.

**Appendix S2**. Characteristics of Linanthus species including number of samples included in analyses, and phenotypic traits.

**Appendix S3**. Location of samples for all the samples used in the ddRADseq phylogenetic and population structure analyses. This sampling covers all the members of Linanthus and includes an average of 7 individuals per species, which is the highest sampling coverage of any previous study. We dispersed the sampling to cover a large part of the species’ ranges. We collected a third of the samples in the field, and sourced two thirds from herbaria across the Western United States.

**Appendix S4**. ipyrad parameters for assembly of RAD data.

**Appendix S5**. AIC scores for models of evolution to use in stochastic character mapping.

**Appendix S6**. Posterior probability density trees of anthocyanin polymorphism evolutionary history with two alternate models of evolution: equal rates and irreversible models.

**Appendix S7**. A distribution of trait changes across 100 stochastic character mapped trees for perenniality, night blooming, and corolla anthocyanin polymorphism.

**Appendix S8**. RAD species trees. A. Species tree with ddRAD data generated with SVDQuartets. B. The same species tree with branch lengths showed inconclusive species relationships due to very short branches. Values represent branch lengths.

**Appendix S9**. TC species trees. A. Phylogenetic tree built with 63 samples across 22 species of Linanthus and 4 outgroups with 219 loci recovered via Target Capture (Angiosperms353) with 100% occupancy generated in ASTRAL. Values represent concordance factors. Most species are recovered as monophyletic. B. Species tree built with 219 loci recovered via Target Capture (Angiosperms353) with 100% occupancy generated in ASTRAL. Values represent concordance factors.

## Notes

https://github.com/ioanaanghel/Linanthus_phylogeny

### Competing Interest Statement

The authors have declared no competing interest.

