## Supplementary figures and images for "Phylogenomics of the North American Desert Radiation *Linanthus* (Polemoniaceae) Reveals Mixed Trait Lability and No Single Geographic Mode of Speciation"

### Appendix S3_RADsamples_locations_wrap

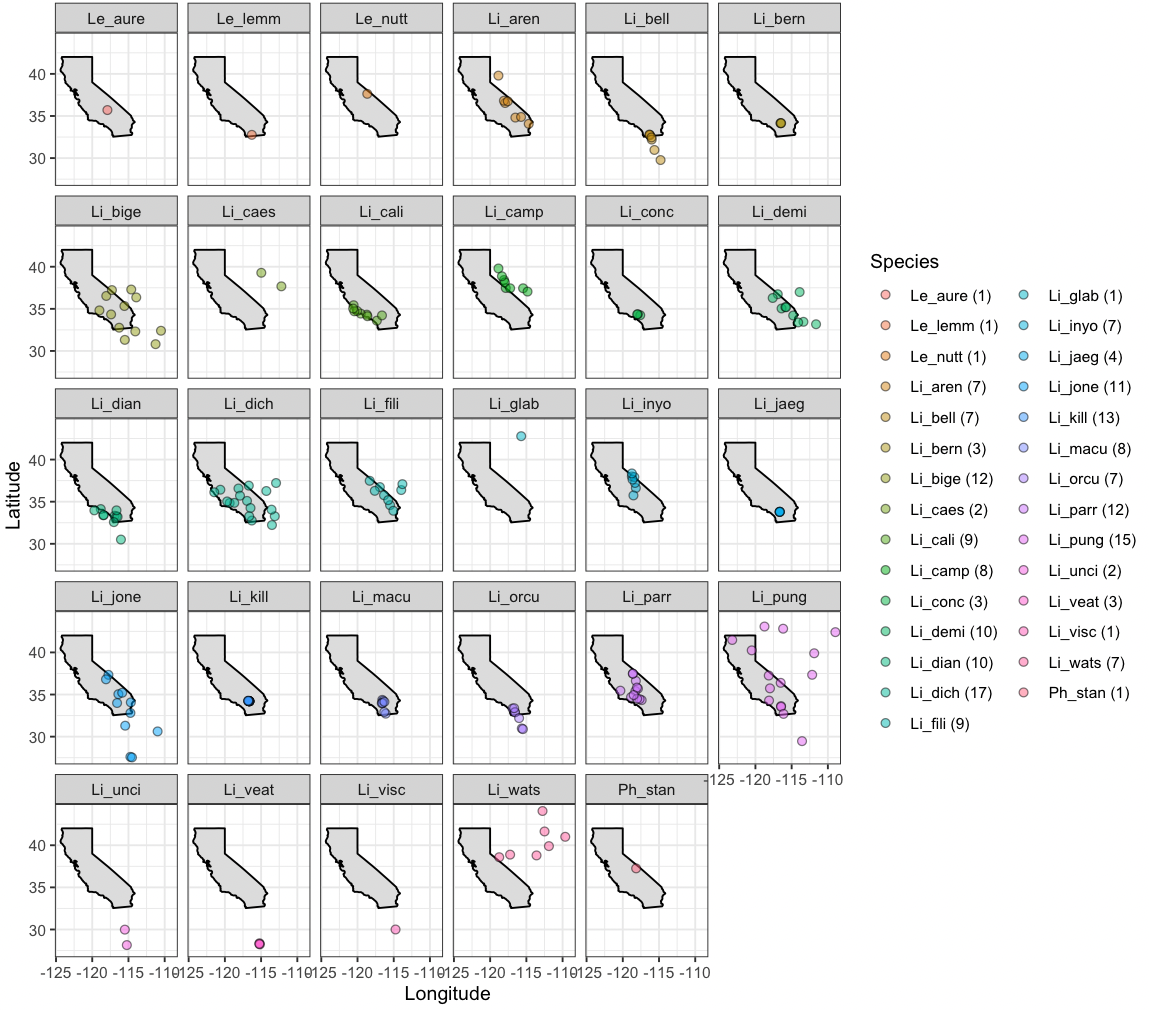

### Appendix S6_Anthocyanins_simmap_postprob_AlternateModels

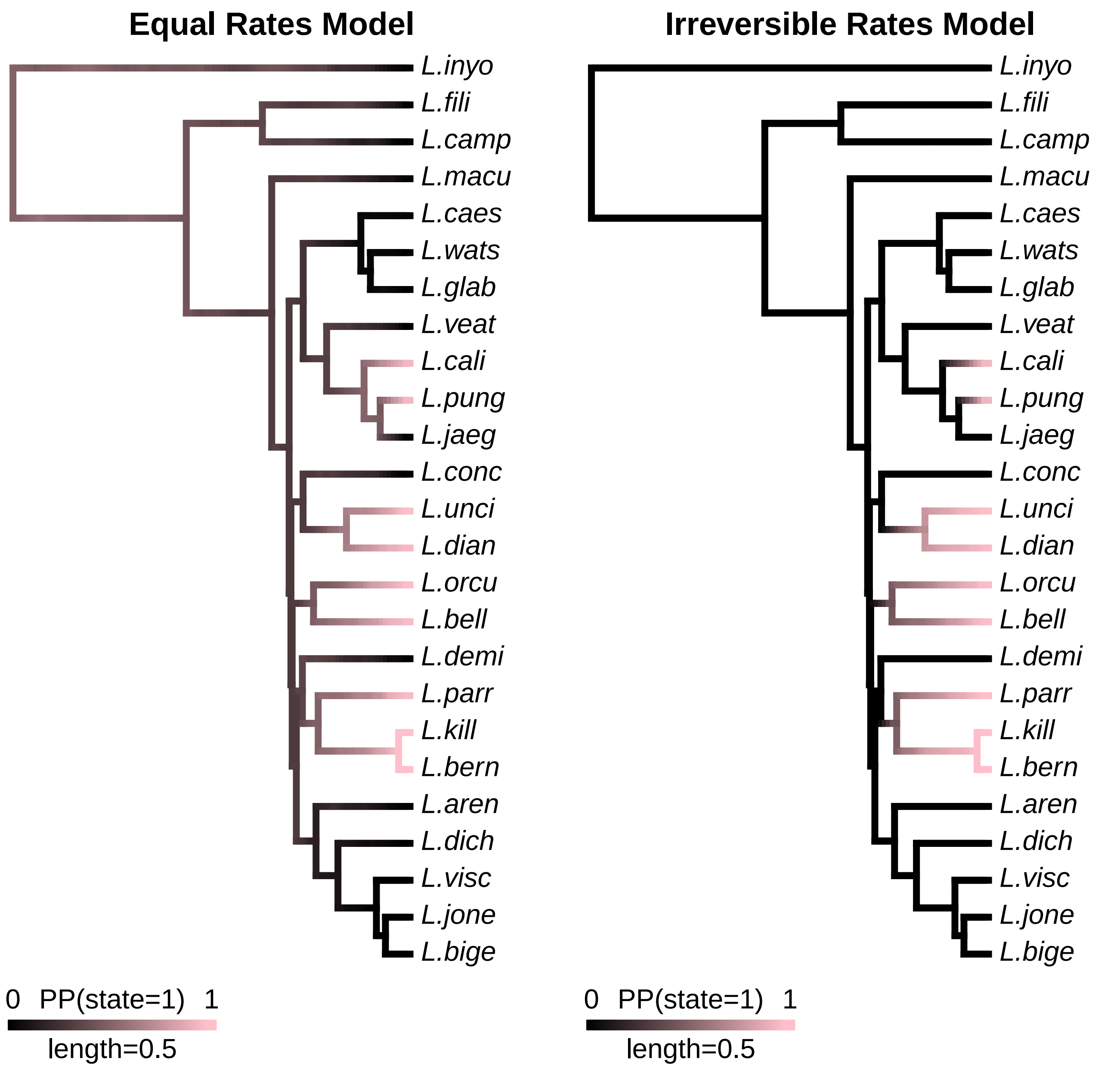

### Appendix S7_Alltraits_simmap_distrib_of_changes

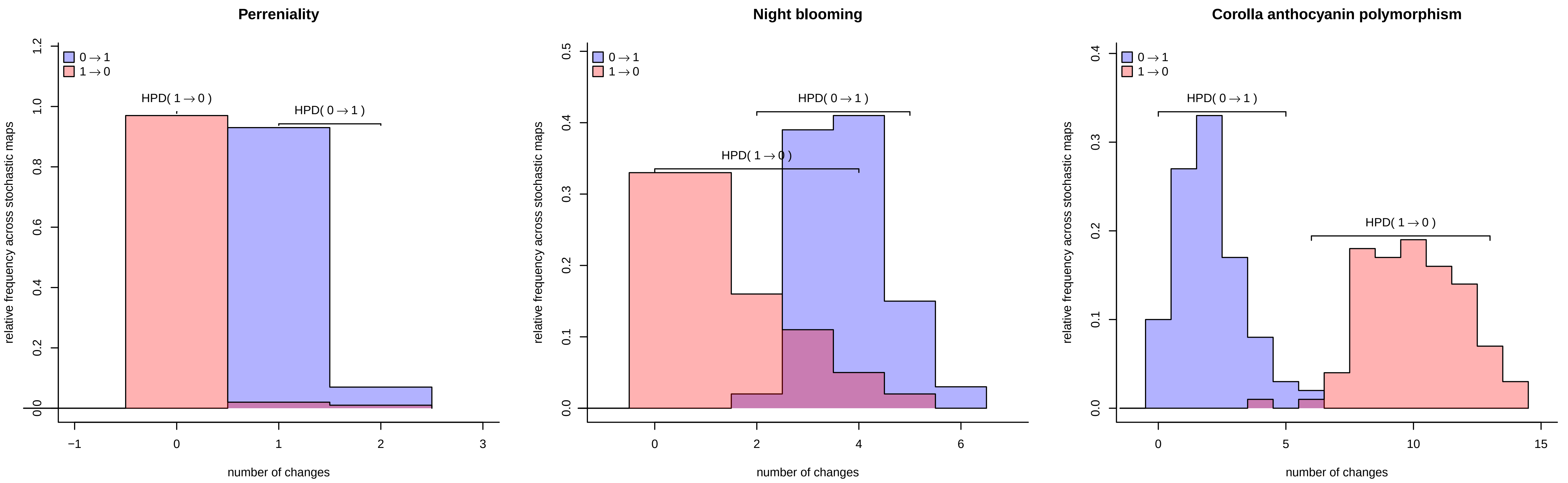

### Appendix S8A_RADSpeciesTree_Li_svdq_min4_ipy_bs100_SpeciesTreenoBL

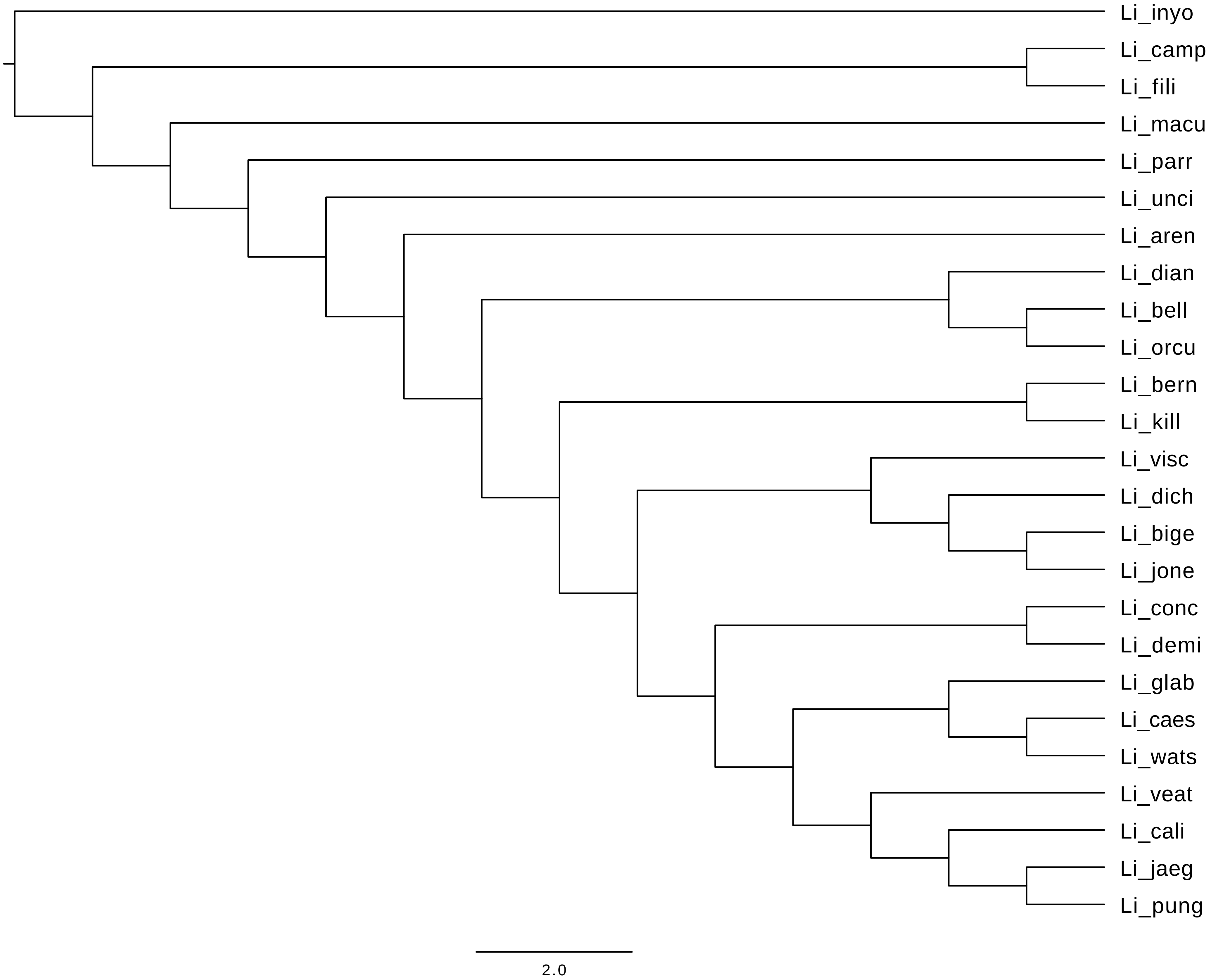

### Appendix S8B_RADSpeciesTree_SVDQ_SpTreewBL_Li_min4_bs100_ipy_SpeciesTree

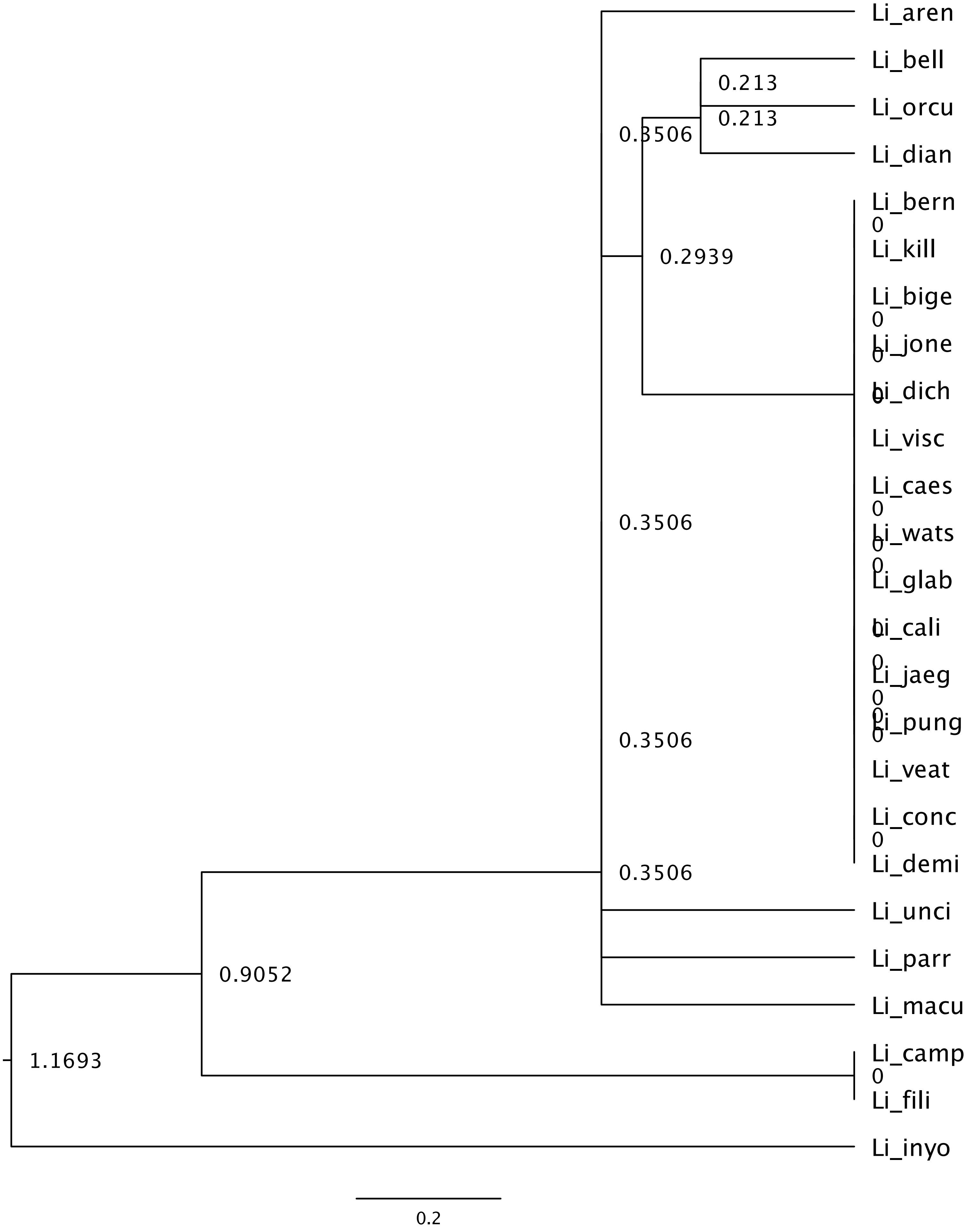

### Appendix S9A_TC_ASTRAL_trimmed_supercontings-0p_219genes_PhylTree

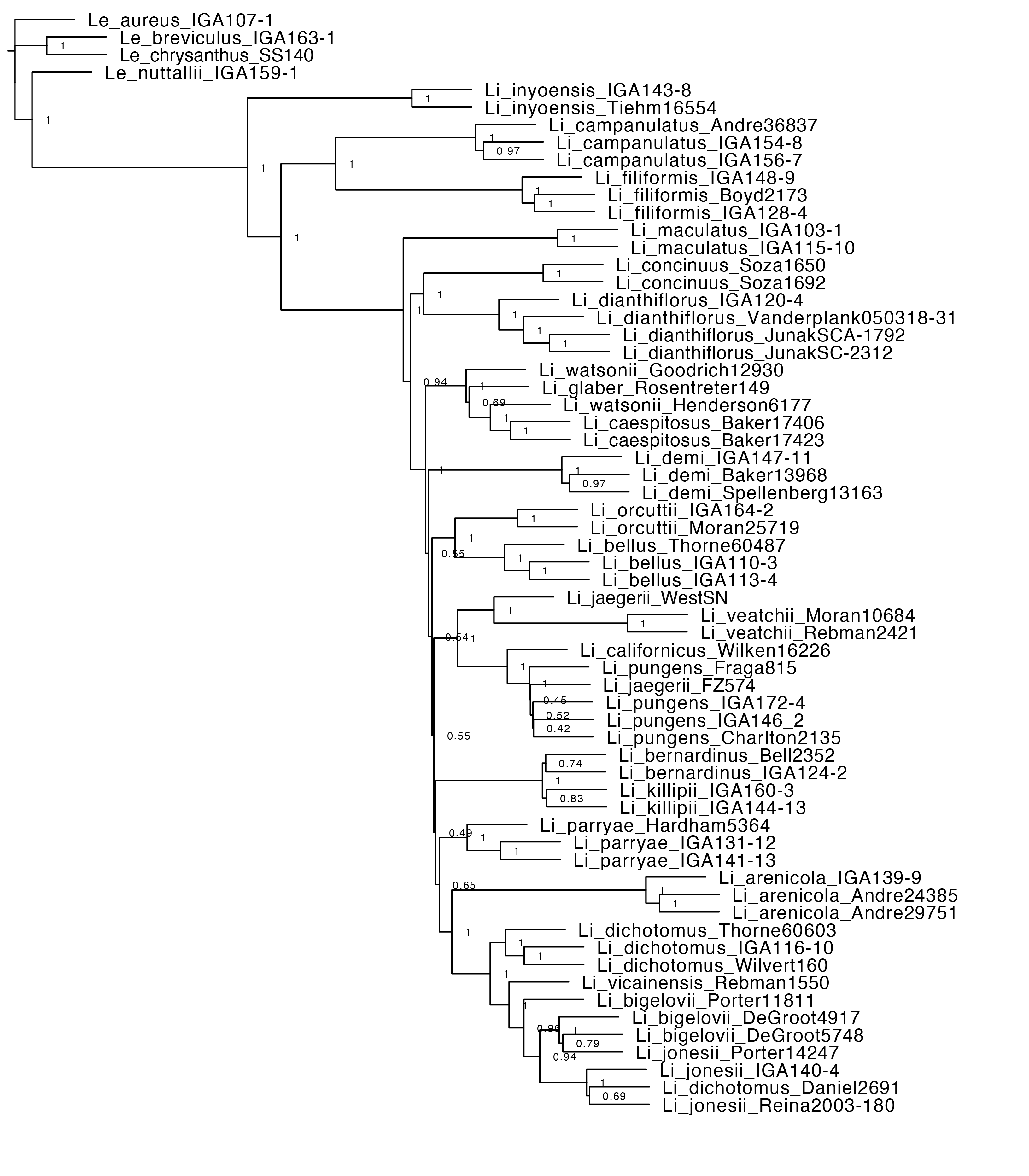

### Appendix S9B_ASTRAL_trimmed_supercontings-0p_219genes_sptree_clades

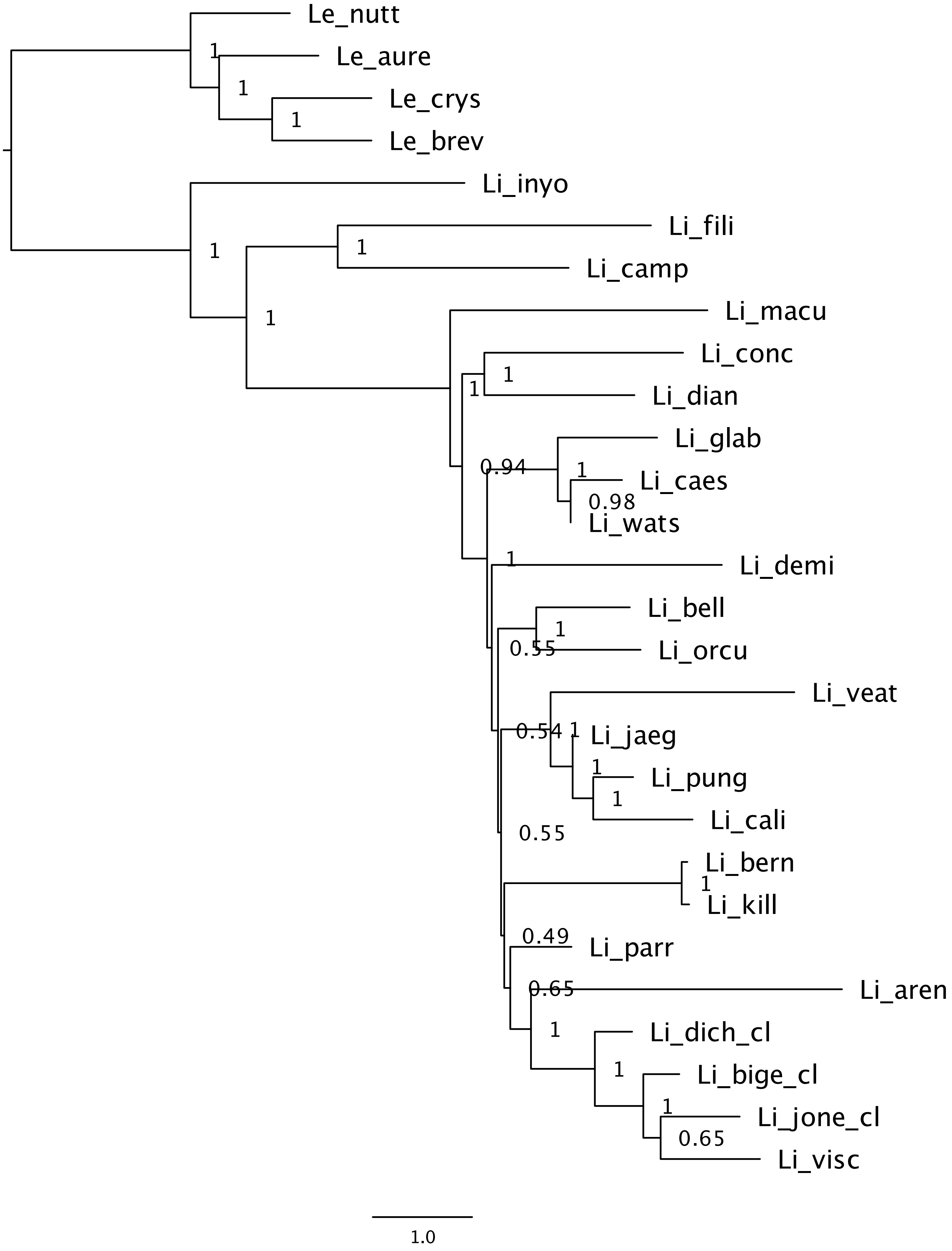
